# Periodic DNA encoding enables error-tolerant multi-class molecular detection by nanopore sequencing

**DOI:** 10.64898/2026.08.17.744993

**Authors:** Mayank Mitram, Manoj Varma

## Abstract

Biomarker analysis requires detecting analyte classes that span nucleic acids, proteins, small molecules, and metabolites, yet testing remains fragmented across target-specific assays and instruments. Here we report a molecular information-transduction strategy that converts target recognition across molecular classes into a common, error-tolerant DNA code readable by nanopore sequencing. Target recognition triggers a hybridization chain reaction that generates concatemers containing periodically repeated 10-nucleotide target-specific barcodes. A matched-filter decoder exploits this periodicity and the linear scaling of read length with match count to reject spurious matches by two to three orders of magnitude. Multi-class detection is demonstrated for a small molecule (ATP), two cardiovascular-associated microRNAs and thrombin in singleplex and multiplexed assays. By separating molecular recognition from sequence readout, this architecture provides a modular framework for converting heterogeneous analytes into a shared, redundancy-encoded signal for high-fidelity molecular sensing.

## Introduction

Disease diagnostics requires the detection of targets ranging from nucleic acids, proteins, small molecules, metabolites, and even whole cells [1–4]. Clinical testing nevertheless remains fragmented into target-specific assays and instruments, for example, the polymerase chain reaction (PCR) is used for genetic markers and immunoassays for protein markers. This fragmentation increases cost and complexity, limits multiplexing and scalability, and impedes access to diagnostic technology for a substantial fraction of the global population [5,6]. A single device and workflow capable of detecting targets across diverse molecular classes could transform the practice of disease diagnostics. Such a multi-class approach faces considerable challenges including cross-reactivity, wide variation in detection limits and dynamic range required for different targets, and differences in surface chemistry and sample-preparation requirements between target classes [6, 7]. Given these constraints, a practical route toward consolidation is a modular architecture in which the presence of any target molecule, whether DNA, RNA, protein, or small molecule, is encoded into a single format that can be read out by a common instrument. DNA barcodes have been employed effectively for highly multiplexed detection [8–10] and more recently nanopores have also been explored for this purpose [11,12]. Sequencing of barcodes is often used as a readout method in these approaches [9,10] and nanopores provide a natural transducer for direct sequencing based multiplex detection.

In a pioneering study, Koch et al. used nanopore sequencing to simultaneously detect microRNAs, proteins, and neurotransmitters [13]. In their assay, an engineered DNA construct is sequenced while binding of the target to a cognate recognition segment produces a characteristic stalling signature during nanopore translocation. Target identity is then recovered by associating this kinetic signature with the corresponding barcoded construct. Although this approach represents an important step toward a consolidated and highly multiplexable workflow, the assay architecture lacks built-in mechanisms for error suppression at the sequence level. The probability of correctly identifying a barcode is consequently limited by the per-read sequencing accuracy. Sequencing errors cause a substantial fraction of reads to be rejected because of poor alignment to the reference barcodes. In the work of Koch et al. [13], approximately 73% of reads were discarded owing to poor alignment quality, reducing process efficiency. Moreover, in a diagnostic setting, a relevant figure of merit is the positive predictive value (PPV). Even a low absolute false-positive rate can appreciably erode the PPV when true-positive events are rare, i.e. for diseases with low prevalence [14]. The absence of an intrinsic error-suppression mechanism is reflected in a false-positive burden of approximately 33% in the work of Koch et al. (0.95% false positives relative to 2.95% true positives) [13].

The problem of recovering a message reliably in the presence of channel noise is foundational in communication engineering, where robustness is achieved by embedding information in a structured codeword that supports reliable detection and decoding [15]. Inspired by this principle, the present work introduces a molecular-to-DNA information-transduction strategy in which aptamer-based target recognition triggers a hybridization chain reaction (HCR) [16] to synthesize a reporter consisting of a periodically repeating barcode, that is, a constrained and non-trivial information structure rather than a single isolated tag. This periodic structure enables a matched-filter-like periodicity score that rejects spurious matches and supports consensus interpretation directly from standard base-called reads, driving the false-positive rates toward zero and the PPV toward unity even for barcodes as short as 10 base pairs (bp). Importantly, the approach is PCR-free and does not require surface immobilization [17, 18] or chemical modification of the biological nanopore [11]. In essence, the strategy recasts molecular diagnostics as an information-transfer problem in which biochemical recognition writes a redundant, noise-tolerant code and nanopore sequencing serves as a universal receiver. The present study establishes this principle of detection across three different target classes. Robust operation in complex biological matrices, and accurate target concentration determination from sequence data are identified as principal directions for future research.

### Detection Strategy

The high-fidelity multiplexed detection strategy combines the hybridization chain reaction (HCR) [16] with nanopore sequencing. For each target, a set of three barcoded hairpins is incubated with the sample, and the products are sequenced on either the PromethION or the MinION platform (Oxford Nanopore Technologies) [19]. The design of the HCR mechanism is shown in Figure 1. Each hairpin set comprises an initiator hairpin (I), a first hairpin (H1), and a second hairpin (H2), built from four oligonucleotide sequence blocks denoted *A*, *B_c_*, *B*_0_ and *C*, and their respective complementary sequences 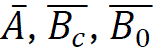 and 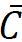. The *B_c_* block is a 10-nucleotide long barcode sequence that is unique to each target. The *C* block likewise differs between targets, whereas the *A* and *B*_0_ blocks are common to all targets. The initiator hairpin (I) additionally carries an aptamer block *R* that binds the target analyte.

Four targets were used in this work, selected to span three important target classes relevant to cardiovascular disease; adenosine triphosphate (ATP), a small-molecule metabolite that reports on cellular energy status and serves as a representative small-molecule analyte; two circulating microRNAs, hsa-mir-1306-5p (AMI) and hsa-miR-18a-5p (CAD), which are associated with acute myocardial infarction and coronary artery disease, respectively [13], and which exemplify nucleic-acid biomarkers detectable in serum and Thrombin (THR), a coagulation protease implicated in thrombus formation [20], which serves as a representative protein analyte. This panel was chosen to demonstrate detection across small-molecule, nucleic-acid, and protein target classes within a single workflow. The specific oligonucleotide and hairpin sequences are provided in Supplementary Information (SI) Table S1. The detailed experimental methodology is provided in SI section 3.

The hairpin sets for all targets are pooled and incubated with the sample. If a target is present, it binds the aptamer block *R* of the corresponding initiator hairpin, triggering a strand-displacement reaction (SDR) in which six nucleotides at the 5ʹ end of the initiator form a single-stranded overhang. The resulting initiator-target complex presents a single-stranded 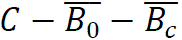 segment toward the 5ʹ end, which undergoes a further SDR with the complementary 3ʹ sticky end of hairpin H2, opening H2 in turn. The cascade of SDRs depicted in Figure 1 propagates through alternating incorporation of H1 and H2, growing a concatemer in which the barcode block *B_c_* is repeated periodically. Each productive recognition event is thereby transduced into a long DNA molecule carrying many copies of the target-specific barcode at a fixed designed spacing.

### Decoding framework

Before presenting the individual assays, the common analysis pipeline and the signatures that distinguish a specific target from non-specific background are defined here. These definitions apply unchanged to every experiment reported below. For each replicate, the sequencing output was divided into chunks of 10 MB. Reads were retained for analysis only if their length was below 2 kb, an upper cutoff that removes contamination from sources unrelated to HCR amplification, whose products do not exceed 1-2 kb (See SI Figure S16, and Reference 16). Across all experiments the great majority of reads satisfied this criterion, indicating good sample quality (for example, SI Figure S1 and S4). Retained reads are referred to as good reads. Each good read was then searched for the presence of each target barcode, and the number of barcode matches in the read, and the interval (in number of nucleotides) between successive matches were recorded. Successful HCR amplification of a target that is present in the sample produces a deterministic concatemer and therefore three concurrent signatures: (i) a barcode interval tightly clustered around the designed value of 48 nt; (ii) a read length that increases linearly with the number of matches; and (iii) a number of matches that can reach high values (for e.g., more than 10). By contrast, barcodes of absent targets are encountered only through random sequence coincidences, whose probability rises with read length. Such spurious matches are characterized by broadly distributed and typically large intervals, no linear relationship between read length and match count, and a small number of matches (typically less than four to six). These contrasting behaviours form the basis of the decoder. Because the hairpin sets for all targets are co-incubated in every reaction, the barcodes of the targets absent from a given sample serve as internal negative controls, sampled under reaction and sequencing conditions identical to those of the cognate signal. This within-sample comparison provides a stringent assessment of detection specificity.

To suppress spurious matches, a matched-filter-like criterion was applied [See SI section 4 for additional details]. A read is counted toward target *i* only if both of the following conditions hold: the measured mean barcode interval *λ_read_* lies within a window *w_λ_* of the designed period, i.e., |*λ_read_* − *λ_des_*| < *w_λ_*; and the read length is consistent with the expected linear growth, (*L_read_*(*m*) − *L*_0_ − *λ_des_m_read_*) < *w_L_*, where *L*_0_ + *λ_des_m_read_* is the expected length of a read containing *m_read_* barcode matches, *λ_des_* is the designed period (48 nt), and *L*_0_ is the pre-barcode length fixed by the design. These filters are applied for all the targets with *i* = 1–4 indexing ATP, miR-AMI, miR-CAD, and THR targets. For each target *i*, *m_read_*(*i*) is the number of matches of the 10-bp barcode *B_c_*(*i*) corresponding to target *i*. The thresholds *w_λ_*and *w_L_* were chosen a-priory and held fixed across all experiments reported in this article. The thresholds chosen for analysis were 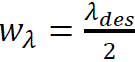 and *w_L_* = *mw_λ_*. Applying these criteria yields *R_i_*(*m*), the number of reads with *m* matches for target *i*; dividing by the total number of reads gives the fractional *R_i_*(*m*).

Two properties of *R_i_*(*m*) are general. Firstly, *R_i_*(*m*) decreases approximately exponentially with increasing *m* for both specific and non-specific targets, the distinction being that a target that is present in the sample yields a markedly higher value of *R_i_*(*m*) at large *m*, for e.g. *m* > 10 [see theoretical model described in SI section 5]. Second, the periodicity and read-length filters together reject spurious matches by two to three orders of magnitude (see for example, SI figure S3 and S6). These observations motivate two complementary visualizations used in this manuscript; a plot of read count against number of matches, shown before filtering, after the periodicity filter, and after both filters, and a two-dimensional map of log fractional *R_i_*(*m*) across all targets, in which the target(s) present produces a pattern that is clearly distinct from the non-specific background. Unless stated otherwise, pooled data across replicates are shown in the main figures and per-replicate results are provided in the Supplementary Information (SI) document.

## Results

### Singleplex detection of ATP

To establish feasibility, the ATP detection experiment of the original HCR study [16] was first reproduced. The pooled hairpin sets for all four targets, each at 1 µM, were incubated with 1 mM ATP, and the mixture was sequenced. The experiment was performed in six replicates (E1-E6). Each replicate yielded between approximately 5,000 and 13,000 reads per chunk (SI Figure S1A). A high fraction of reads passed the 2-kb length filter (SI Figure S1B). For the replicate with the fewest good reads (E1; 95% good reads), the median read length was approximately 400 bp, with most reads well below 2 kb (SI Figure S1C, S1D). The error bars in all figures denote the 25th and 75th percentiles. The asymmetry of the read-length distribution about the median is consistent with a potential minor contribution from contaminating DNA or chimeric molecules, which get removed by the read length filter. Read-length distributions for the remaining replicates are shown in SI Figure S1.

Searching the filtered reads for each barcode revealed the expected specific signatures for ATP and their absence for the other three targets (Figure 2). The ATP barcode interval clustered tightly around the designed value of 48 bp, whereas the intervals for miR-AMI, miR-CAD, and THR were broadly distributed and centred near 2,000 bp, consistent with random coincidences in longer reads (Figure 2A, 2B). Read length increased linearly with match count for ATP but not for the other targets (Figure 2C, 2D), and the ATP barcode reached as many as 14 matches per read, whereas the non-specific barcodes did not exceed four or five (Figure 2D). Each of these features is a direct consequence of deterministic HCR amplification of the ATP hairpin set and confirms specific single-plex detection. Application of the periodicity and read-length filters suppressed spurious matches by approximately two to three orders of magnitude (SI Figure S3). For example, without filtering approximately 50 reads exhibited five miR-CAD matches, *R_CAD_*(5). After the application of the periodicity and read length filters *R_CAD_*(5) fell to zero, whereas *R_ATP_*(5) remained near 10,000. The rule of three [21] gives an approximate 95% upper confidence bound of 3/N on the per-read false-positive probability, where N is the number of reads. The false-positive rate is bounded above this value, and the positive predictive value (PPV) can be made arbitrarily close to unity by increasing N. The two-dimensional map of log fractional *R_i_*(*m*) shows the ATP signal being clearly distinct from the other non-specific targets, with progressive improvement as each filter is applied (Figure 3A and 3B). The data shown are pooled across the six replicates. The per-replicate results, together with the corresponding rejection fractions, are provided in SI Figure S2 and S3, respectively. Finally, we investigated if there was any bias in the readout between the top and bottom strands of the double stranded HCR product (Figure 1) by inspecting the sequences before (prefix) and after (suffix) the barcode region. This barcode flanking region analysis is presented in detail in SI section 4 and shows that the top and bottom strands are equally represented in the sequence data (SI figures S17 – S19).

**Figure 1.**
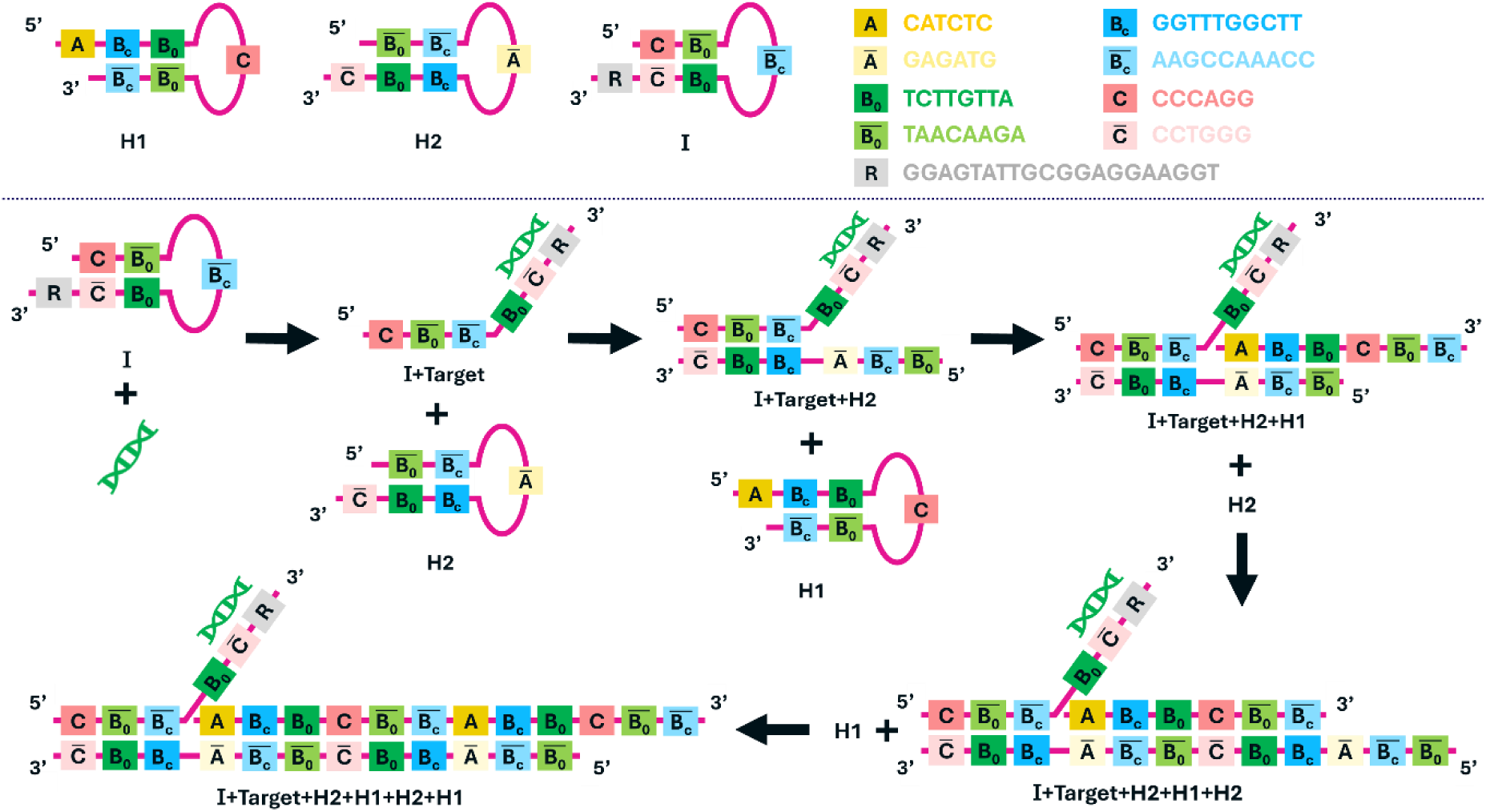
Target-to-sequence transduction strategy. Schematic illustration of the encoding mechanism. The target molecule opens initiator hairpin I, which exposes a single-stranded segment that opens hairpin H2 to form a partial duplex with I. The remaining single-stranded segment of H2 then opens hairpin H1, forming a partial duplex with H2. Repetition of this strand-displacement cascade extends the concatemer, producing periodic repeats of the target-specific barcode *B_c_*.

**Figure 2.**
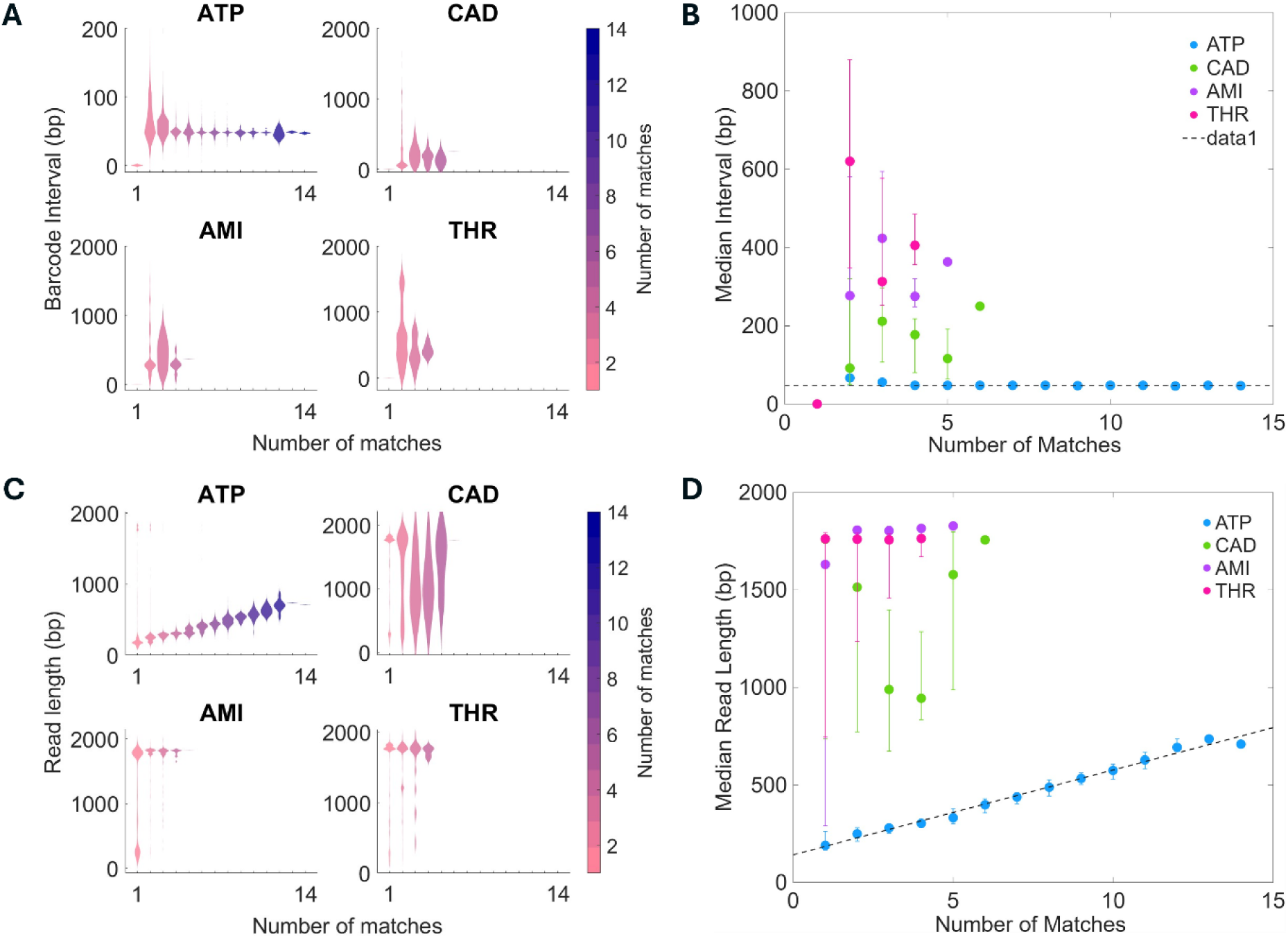
Barcode-interval and read-length signatures for the ATP singleplex assay. Barcode interval and read length as a function of the number of matches for the specific target (ATP) and the non-specific targets (miR-AMI, miR-CAD, THR), showing tight clustering near 48 bp (A, and B) and linear length scaling only for ATP (C, and D).

**Figure 3.**
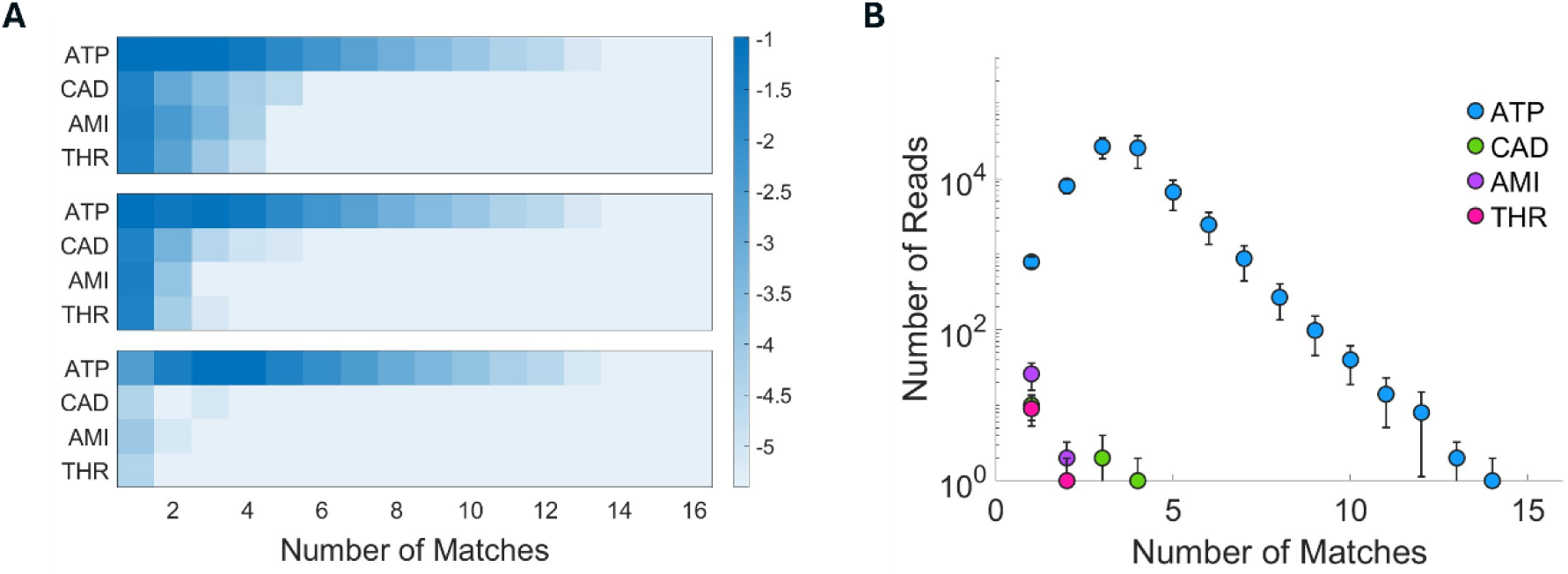
Filtered barcode-match statistics for the ATP singleplex assay. (A) Two-dimensional map of log fractional *R_i_*(*m*) across all four targets before filtering (top row), after the periodicity filter (middle row), and after both filters (bottom row). The filtering significantly suppresses signal from non-specific targets while retaining the specific target signal. (B) Read count as a function of the number of matches after application of both filters.

### Multiplexed detection

Two multiplexed configurations were tested. In the first, the pooled four-target hairpin sets were incubated with miR-AMI and miR-CAD, each at 200 nM, in duplicate. In the second, the pooled sets were incubated with miR-AMI and miR-CAD at 200 nM and THR at 1 U/µL, in four replicates. As before, the sequencing output was divided into 10-MB chunks. To test whether the match count could exceed the maximum of 14 observed in the ATP assay, the multiplexed mixtures were sequenced to greater depth. In both configurations the targets that were present in each sample reproduced the two specific signatures associated with successful decoding of specific target binding, i.e., tight clustering of gaps between successive matches of the target barcode around the designed value of 48 bp, and the linear relationship between read length and number of matches. The targets that were absent in the sample did not exhibit these features. The quantitative details of these experiments are reported below.

For the miR-AMI/miR-CAD duplex, the two replicates yielded 2.64 million and 1.71 million reads (E1 and E2), corresponding to roughly 7,500-12,000 reads per chunk (SI Figure S4A); more than 99% passed the length filter (SI Figure S4B). Median read lengths were approximately 200 bp (E1) and 400 bp (E2), with E2 showing greater chunk-to-chunk variability (SI Figure S4C, S4D). Both targets present (miR-AMI and miR-CAD) exhibited intervals clustered near 48 bp and linear length scaling, whereas ATP and THR did not (Figure 4A-D). The miR-AMI and miR-CAD barcodes reached as many as 40 and 30 matches per read, respectively, compared with maxima of about six (ATP) and three (THR). After matched filtering, the two-dimensional map and the read-count distributions cleanly isolated miR-AMI and miR-CAD (Figure 5A, 5B; SI Figure S6). These results are pooled across the two replicates with the per-replicate results shown in SI Figure S5 (barcode-interval and read-length) and S7 (two-dimensional map, read-count distributions, and rejection ratios).

**Figure 4.**
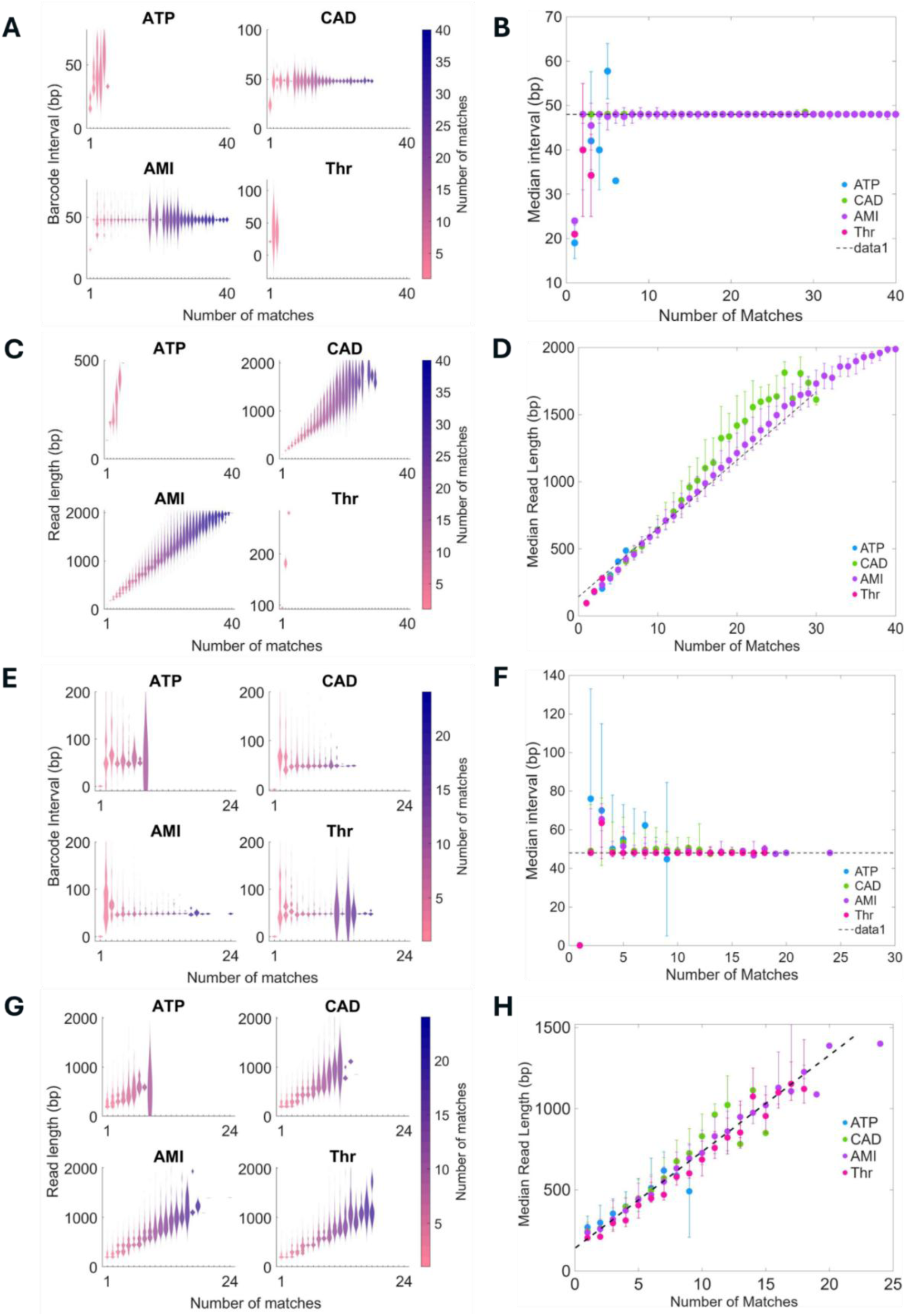
Barcode-interval and read-length signatures for the multiplexed assays. Barcode interval and read length as a function of the number of matches for the miR-AMI/miR-CAD duplex (A-D) and the miR-AMI/miR-CAD/THR triplex (E-H). Present targets show clustering near 48 bp (A, B, E, F) and linear length scaling (C, D, G H) while absent targets do not.

**Figure 5.**
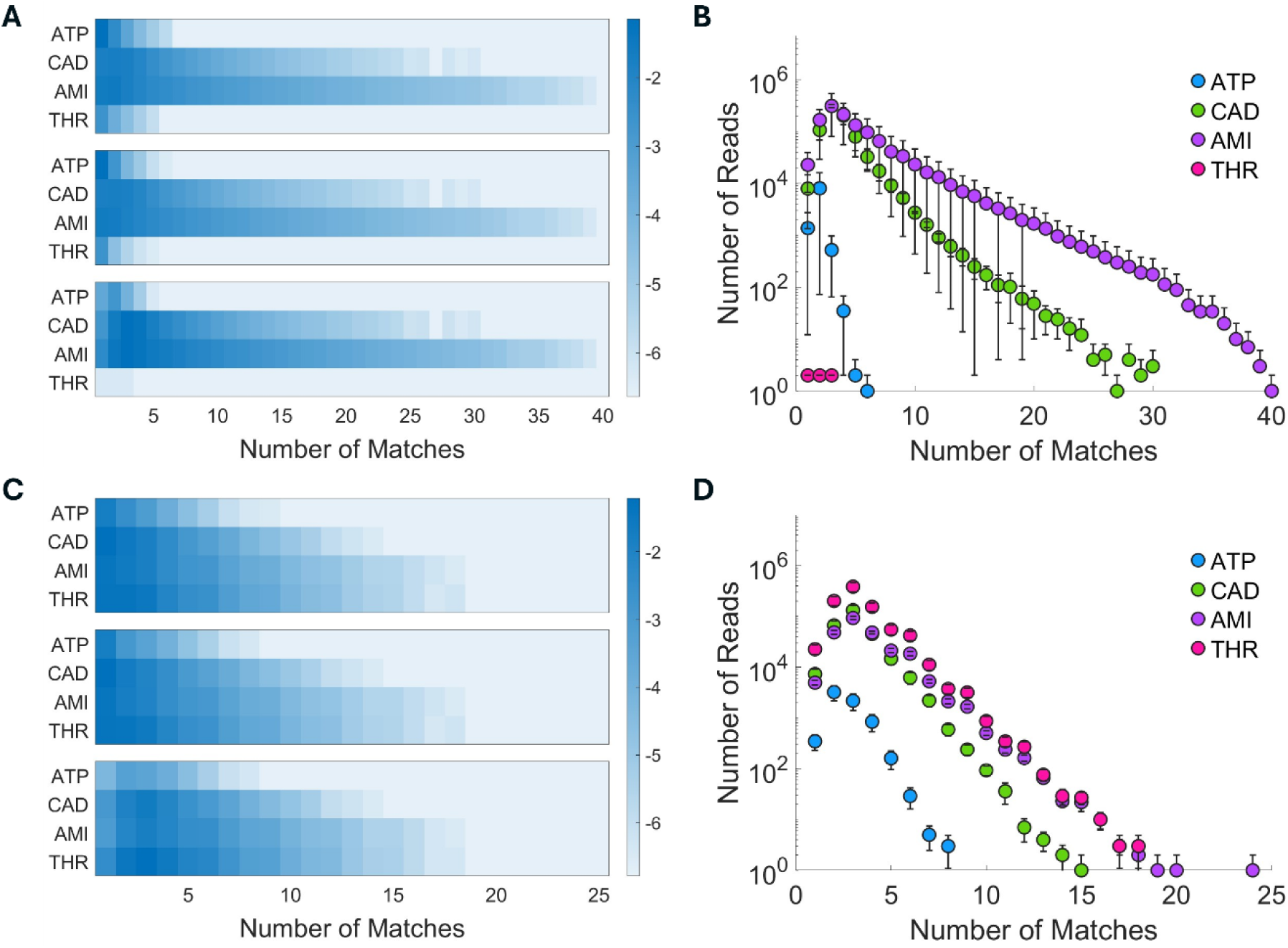
Filtered barcode-match statistics for the multiplexed assays. Two-dimensional maps of log fractional *R_i_*(*m*) and read-count distributions for the miR-AMI/miR-CAD duplex (A, B) and the miR-AMI/miR-CAD/THR triplex (C, D), each showing distinct patterns for the present targets after filtering.

For the miR-AMI/miR-CAD/THR triplex, the four replicates yielded approximately 9,000-11,500 reads per chunk (SI Figure S8A), of which more than 99% passed the length filter (SI Figure S8B). The median read length was approximately 250 bp and was reproducible across replicates (SI Figure S8C, S8D). All three present targets showed intervals clustered near 48 bp and linear length scaling, whereas the absent target ATP did not (Figure 4E-H). The miR-AMI, miR-CAD, and THR barcodes reached as many as 24, 16, and 19 matches per read, respectively, whereas ATP reached at most seven. The residual ATP signal arises from random matches and was substantially suppressed, though not entirely eliminated, by the matched filtering approach. After matched filtering, the two-dimensional map and read-count distributions isolated the three present targets (Figure 5C, 5D; SI Figure S10). Data are pooled across the four replicates with the per-replicate results shown in SI Figure S9 and S11.

### Cross-platform validation on the MinION

The ATP singleplex and the two multiplexed samples described above were sequenced on the Nanopore PromethION sequencer (PromethION24 and Data Acquisition Unit, Oxford Nanopore Technology, UK) by an external service provider Genotypic [22]. To assess the robustness of the detection strategy across sequencing platforms and assay sites, single-plex detection of approximately 670 nM miR-CAD against a background of the ATP, miR-AMI, and THR hairpin sets was run in-house using the MinION MK1B sequencer (MIN-101B). The PromethION flow cells (FLO-PRO114M) contain up to 12,000 pores whereas the MinION flow cells (FLO-MIN114) contain up to 2,048 pores. However, the nanopore chemistries in both platforms were kept the same (R10 series, V14 kits).

For the miR-CAD singleplex, between approximately 13,000 and 18,000 reads were obtained per chunk (SI Figure S12A), with 98-99% good reads (SI Figure S12B). However one significant difference observed was a reduction in the median read length to approximately 100 bp compared to 250-400 bp for experiments done at the external site (SI Figure S12C, S12D). We believe this reduction is due to variations in the sample processing between the two sites. Despite this difference, the miR-CAD barcode showed the specific interval and linear-scaling signatures corresponding to successful signal decoding. However, the maximum matches observed for miR-CAD was only eight with matches corresponding to the absent targets being fewer than four (Figure 6A-D). Thus, although from the point of discriminating the specific target from the background, the MinION assay was successful, sample processing differences have likely resulted in a diminished performance with respect to producing reads with large number of matches, for example greater than 15. Nevertheless, matched filtering isolated the miR-CAD signal from the background (Figure 6E, 6F; SI Figure S13).

**Figure 6.**
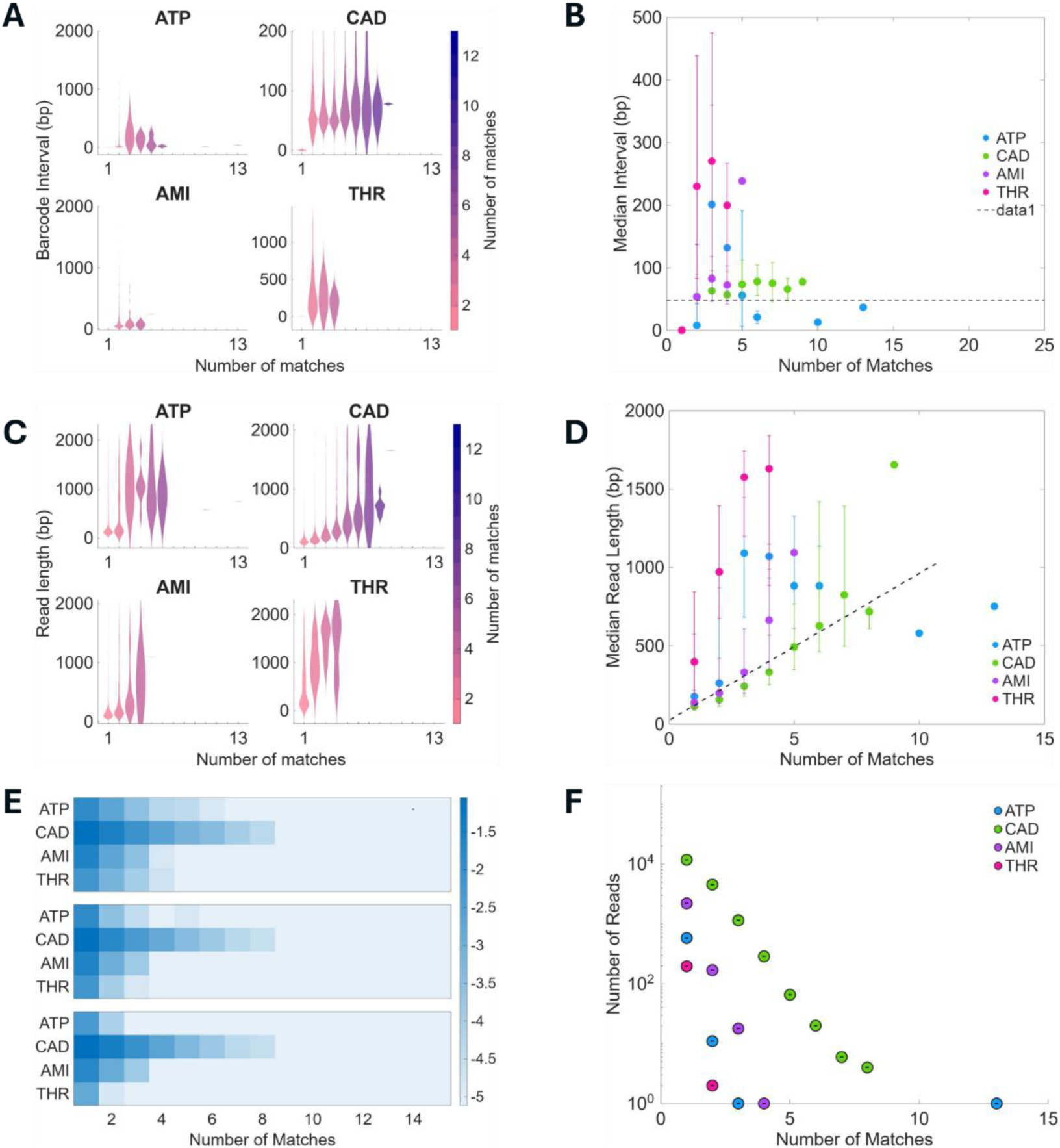
MinION singleplex detection of miR-CAD. Barcode-interval and read-length signatures (A-D) and filtered barcode-match statistics (E, F) for miR-CAD detected against a background of non-cognate hairpin sets.

### Detection in a bacterial genomic-DNA background

The experiments described so far were performed in reaction buffer containing no interfering analytes beyond the pooled hairpin sets, providing well-controlled conditions under which both singleplex and multiplexed detection were demonstrated with high specificity. To assess robustness in a more realistic sample condition, miR-AMI was detected in a background of bacterial whole-genome DNA, since extraneous genomic DNA is a common contaminant in environmental and clinical samples. Genomic DNA was extracted from an *E. coli* K-12 cell lysate and used as the background into which the four HCR hairpin sets and 200 nM miR-AMI were added.

The results of this assay exhibited read statistics that deviated significantly from other experiments. Approximately 1,000 reads per chunk were obtained compared to typical values of 5000-10000 in other experiments (SI Figure S14A). Median read length of approximately 1,200 bp was obtained (SI Figure S14C, S14D), notably longer than in buffer. This could be explained by the presence of extraneous genomic DNA in the sample. Most importantly, the fraction of good reads in almost all the chunks was substantially less than the typical 98%-99% observed in previous experiments (SI Figure S14B). Less than 5% of the reads had the good-read fraction above 90%. Thus, the sequencing-output characteristics of this sample differed substantially from those of the buffer-based assays.

Despite this altered sequence background, the read-length-versus-match-count relationship remained approximately linear for the cognate miR-AMI barcode, indicating that productive miR-AMI-triggered HCR was retained (Figure 7A-D). However, the barcode-interval and read-length signatures were less clearly separated from the non-specific background than in the buffer assays. Thus, the principal effect observed in this experiment was a reduction in the contrast between the specific HCR signal and the background rather than a complete loss of target-triggered HCR. Importantly, application of the periodicity and read-length filters did not suppress the non-specific signals as effectively as in the buffer assays, and residual signals corresponding to the absent ATP and THR targets remained after filtering (Figure 7E, 7F; SI Figure S15). The origin of these signals cannot be uniquely established from the present experiment. One plausible contribution is the increased occurrence of fortuitous matches to the short 10-nt barcodes within the large bacterial genomic-DNA sequence background, including matches introduced or enhanced by sequencing errors. In addition, matrix-dependent non-specific HCR or trace contaminants carried through genomic-DNA purification cannot be excluded. The relative contributions of these mechanisms were not independently quantified in the present study.

**Figure 7.**
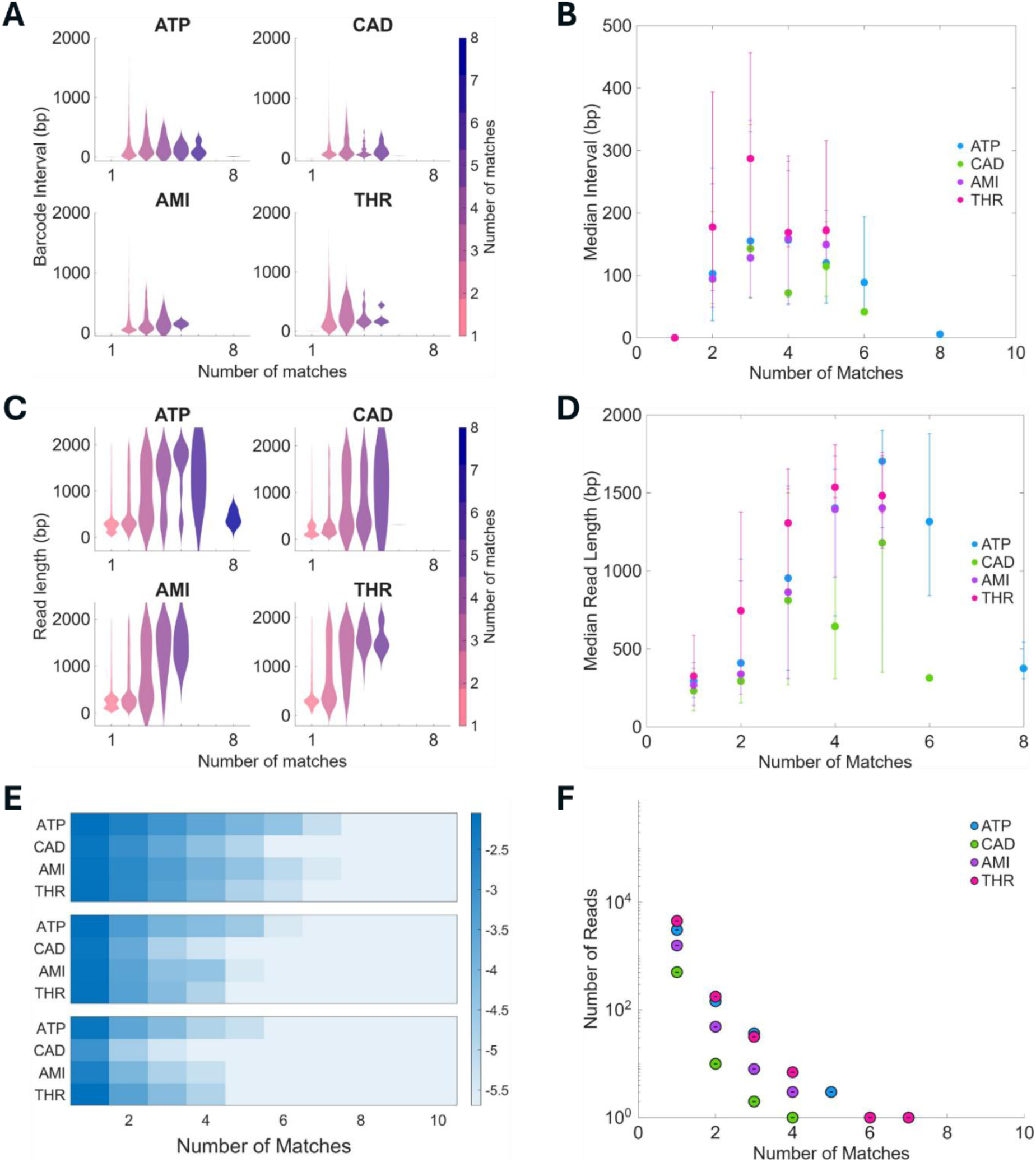
Detection of miR-AMI in a bacterial genomic background (E. coli K-12) Barcode-interval and read-length signatures (A-D) and filtered barcode-match statistics for the 10-bp barcode (E, F). The cognate miR-AMI signal remains detectable, while residual non-specific barcode-associated signals corresponding to ATP and THR are observed.

These findings are preliminary and are included to indicate that the loss of specificity in complex matrices is plausibly remediable rather than fundamental. These results demonstrate that target-triggered HCR remains detectable in a bacterial genomic-DNA background, while also identifying important limitations of the present implementation. Increasing the barcode length, optimizing sample preparation and HCR conditions, and incorporating more error-tolerance in barcode design to improve performance will be the subject of future investigations.

### Concentration determination

Determination of the target concentration in a sample is a critical requirement in most molecular analysis workflows although a binary yes/no result may be satisfactory for a subset of applications. A principal limitation of the detection strategy described in this work is the absence of a simple relationship between target concentration and the barcode-match statistics. To characterise the dependence of the per-read match count on target abundance, a stochastic reaction model was developed, in which an aptamer-bearing initiator is activated by target binding with a dimensionless reaction rate parameter *α*. The activated initiators are subsequently labelled by the stochastic incorporation of barcodes from a finite shared pool, with another dimensionless rate parameter *β*. This model was simulated via tau-leaping method [23] and is described in detail in SI section 5, including the table of parameter values used to obtain the simulation results shown below.

Figure 8A shows the log fractional *R_i_*(*m*) as a function of the normalized target concentration for different values of *α* (denoted in the figure as “activfrac”). The color represents the magnitude of the log fractional *R_i_*(*m*). As we note from the figure, *R_i_*(*m*) is not a monotonic function of the target concentration, in general (Figure 8B). For small values of *m*, for example, *m* = 1, *R_i_*(*m*) is approximately monotonic. However, small values of *m* are susceptible to significant non-specific signal contribution (e.g. Figure 3B). Ideally, one would like to choose *m* > 5 to achieve almost complete suppression of non-specific signal. However, as noted from Figure 8B, target concentration extracted from *R_i_*(*m*) for *m* > 5 is not unique. The dashed lines in Figure 8B indicate examples of values of *R_i_*(*m*) for *m* = 5 and *m* = 10 for which two different target concentrations satisfy the observed data. Such problems apply not only to *R_i_*(*m*), but also to simple composite quantities such as the sum or the area under the curve (AUC) of *R_i_*(*m*) vs *m* (Figure 8C). Here the notation AUC(*k*) refers to summation of *R_i_*(*m*) for *m* ≥ *k*. AUC(1) may seem better suited for concentration estimation from the measured signal due to its approximately monotonic behavior but is more susceptible to non-specific signal due to spurious barcode matches at low *k*. More robust quantities at higher *k*, such as *AUC*(6), unfortunately, cannot be mapped uniquely to a target concentration. Thus, at present the detection strategy described here is primarily applicable for a binary yes/no multiplexed sensing. Further work is required to establish accurate concentration estimation from the sequenced data. Despite these shortcomings, figure 8D shows that the stochastic model provides a close match to the experimental data. The model deviates from experimental observation for small *m*, (e.g. *m* < 3). This is ascribed to the filters rejecting some of the true barcode matches for the specific target. With no filters applied, or with only the periodicity filter applied, the match is close even for small *m* (SI Figure S24). Unfortunately, due to the large uncertainties in connecting the simulation parameters to actual experimental conditions, such apparently close matches can be obtained for other combinations (SI section 5.6 and SI Figure S23) of parameters emphasizing the importance of further studies to connect experimental conditions to models and exploration of data analysis methods to accurately extract the target concentration from sequence data.

**Figure 8.**
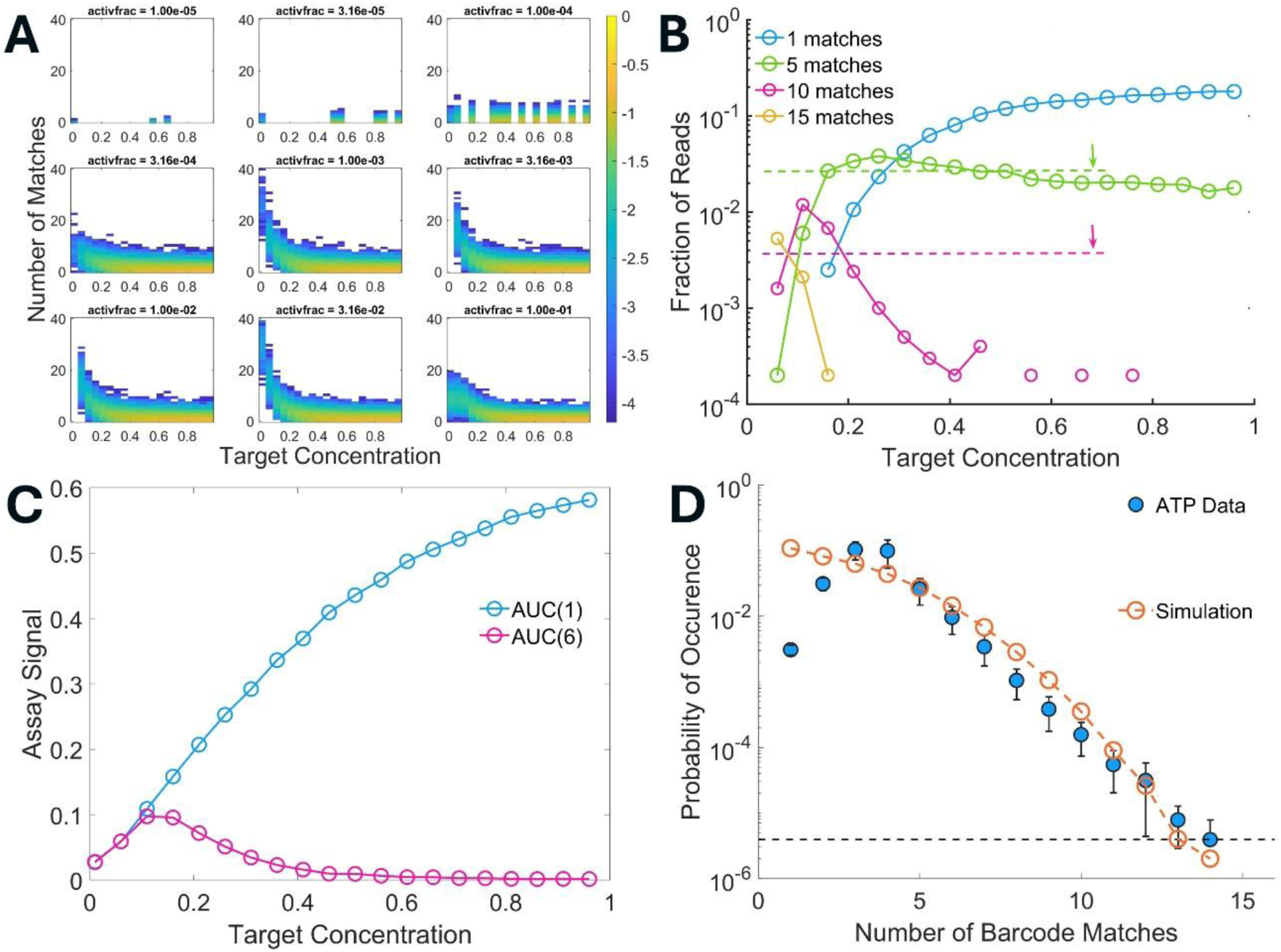
Modelled dependence of barcode-match statistics on target concentration. Tau-leaping simulation of aptamer-triggered HCR, in which initiators are activated at a normalized rate *α* (denoted as “activfrac” in sub-panel titles) and labelled from a finite barcode pool at rate β concentrations are normalized to the initial initiator concentration. (A) Heatmaps of log fractional *R_i_*(*m*) versus match count *m* as a function of normalized target concentration for different *α*. (B) Read fraction versus concentration at fixed *m* exhibiting non-monotonic behavior. (C) Composite signal AUC(*k*) for *k* = 1 and 6, AUC(1) is monotonic but likely to be noise-prone due to spurious barcode matches at low *k*, whereas more robust quantities such as AUC(6) display a non-monotonic behavior with respect to target concentration. (D) Comparison between experimental data and stochastic model for the ATP single-plex assay showing good agreement.

### Conclusions

This work reframes multiplexed molecular detection as an information-transfer problem and demonstrates a corresponding encode-and-decode strategy. Aptamer-triggered HCR transduces a target-recognition event into a long DNA concatemer carrying a periodically repeating, target-specific barcode, and a matched-filter-like decoder exploits the designed periodicity, together with the linear scaling of read length with match count, to separate true signals from the random barcode coincidences that dominate the background. The redundancy written into the reporter allows reliable detection with barcodes as short as 10 bp while suppressing spurious matches by two to three orders of magnitude, and the scheme operates directly on standard base-called reads without PCR, surface immobilization, or modification of the nanopore. Detection was demonstrated for a four-member panel spanning a small molecule, two microRNAs, and a protein, in single-plex and multiplexed formats, on both the PromethION and the MinION, and in a bacterial whole-genome background. We also note that limit of detection could be improved by incorporating amplification steps.

Several limitations constrain the scope of the present study and indicate directions for further work. The panel comprises four targets and scaling to the dozens of analytes required for clinical panels will depend principally on the availability of suitable aptamers and on the design of mutually orthogonal barcodes. Concentration estimation from sequencing data remains an open problem. Finally, although multi-class detection is demonstrated here in controlled backgrounds, operation in complex biological matrices remains to be established. The reduced performance observed in a bacterial-genomic background is offered as preliminary evidence that such matrices are challenging but not necessarily prohibitive, with mitigating sample-preparation and assay design protocols left to future work. Increasing the barcode length offers a complementary route to improved specificity in such matrices, since longer barcodes reduce the probability of random sequence coincidences within large genomic backgrounds. Notwithstanding these open questions, the results establish a biochemical recognition strategy that writes a target-specific DNA code that nanopore sequencing reads as a universal receiver, providing a route toward consolidated, high-fidelity, multi-class molecular diagnostics.

## Acknowledgements

We acknowledge funding support from Blockchain for Impact (BFI) under the grant BFI/IISc/4/25. We also thank the staff members at Genotypic Technologies Pvt. Ltd. Bangalore for technical support.

## Supplementary Information

### 1. Table of the HCR probe sequences used in this work

**Supplementary Table 1:**
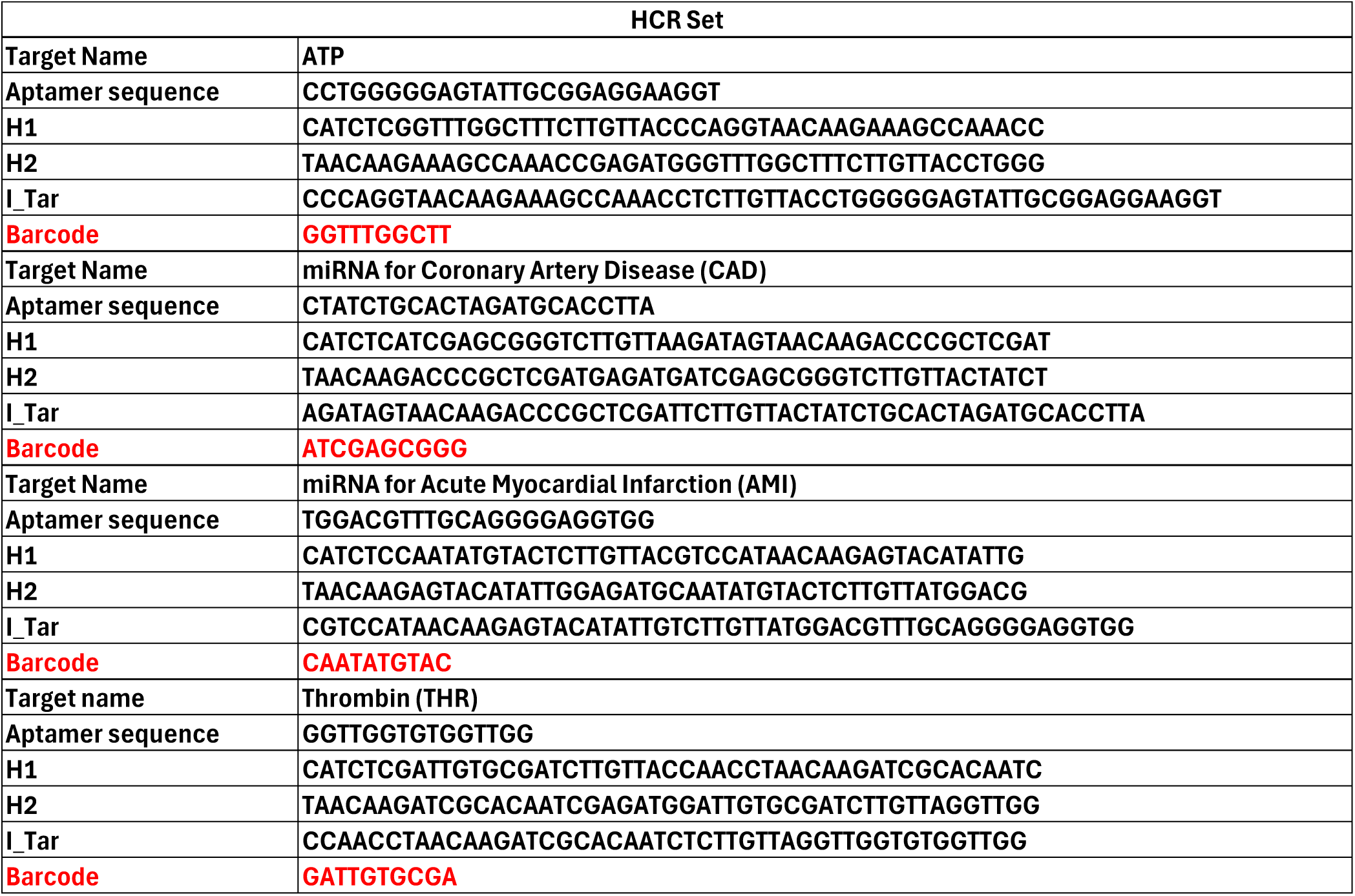
Hairpin sequences for the targets used in this work.

**Supplementary Table 2:**
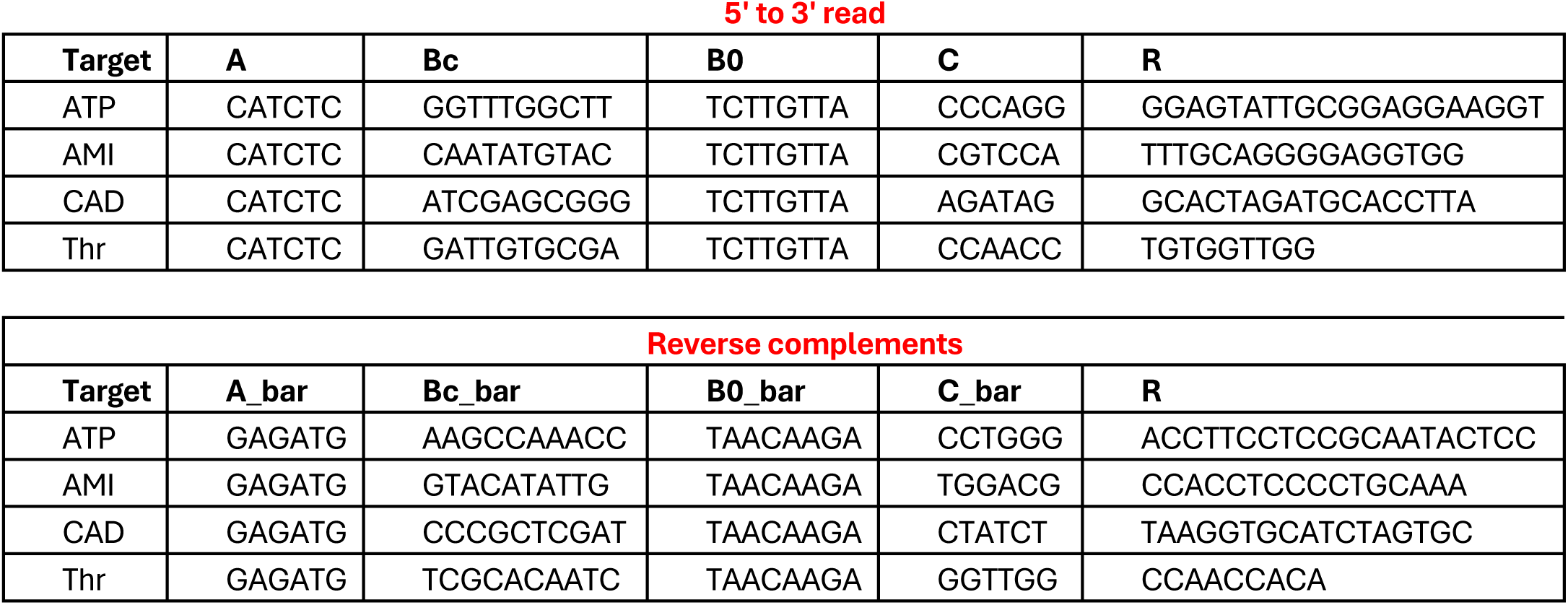
Sequences corresponding to the blocks forming the HCR hairpin sets. The block names refer to the schematic shown in Figure 1 of the main text.

### 2. Supplementary Figures

Throughout this section, “good reads” denotes reads passing the 2-kb length filter, the barcode interval is the spacing in nucleotides between successive barcode matches, and “specific” and “non-specific” targets refer respectively to analytes present in, and absent from, a given reaction. In all read-length distributions, the upper and lower error bars denote the 75^th^ and 25^th^ percentiles, respectively. The same panel layout is used for each experiment so that results may be compared directly across assays.

#### ATP singleplex assay

**Supplementary Fig. S1:**
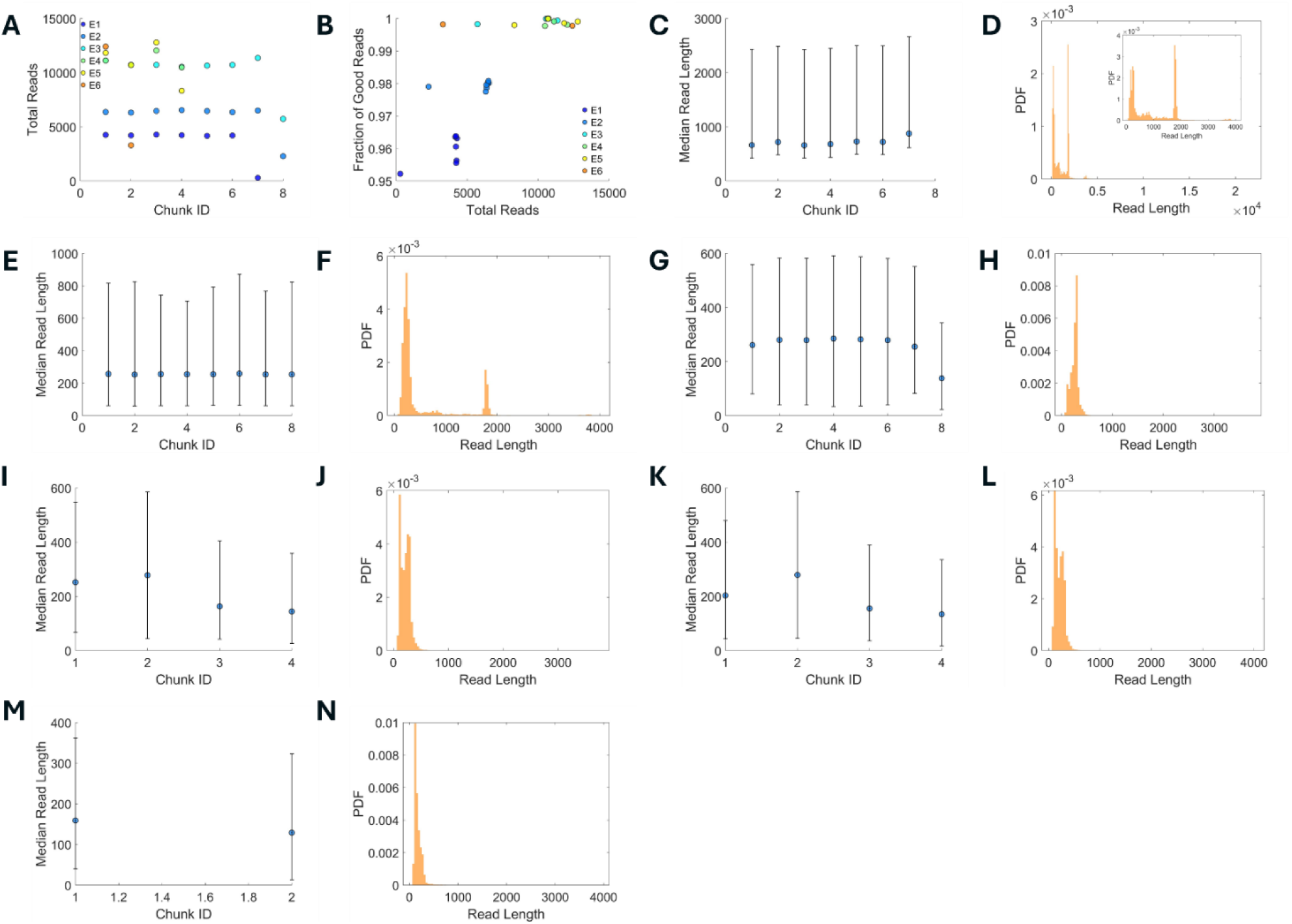
Read-length statistics for the ATP singleplex replicates. (A) Total reads per chunk for all chunks across the six replicates. (B) Fraction of good reads for each replicate. (C, E, G, I, K, M) Median read length per chunk for replicates E1-E6. (D, F, H, J, L, N) Probability density of read length for replicates E1-E6.

**Supplementary Fig. S2:**
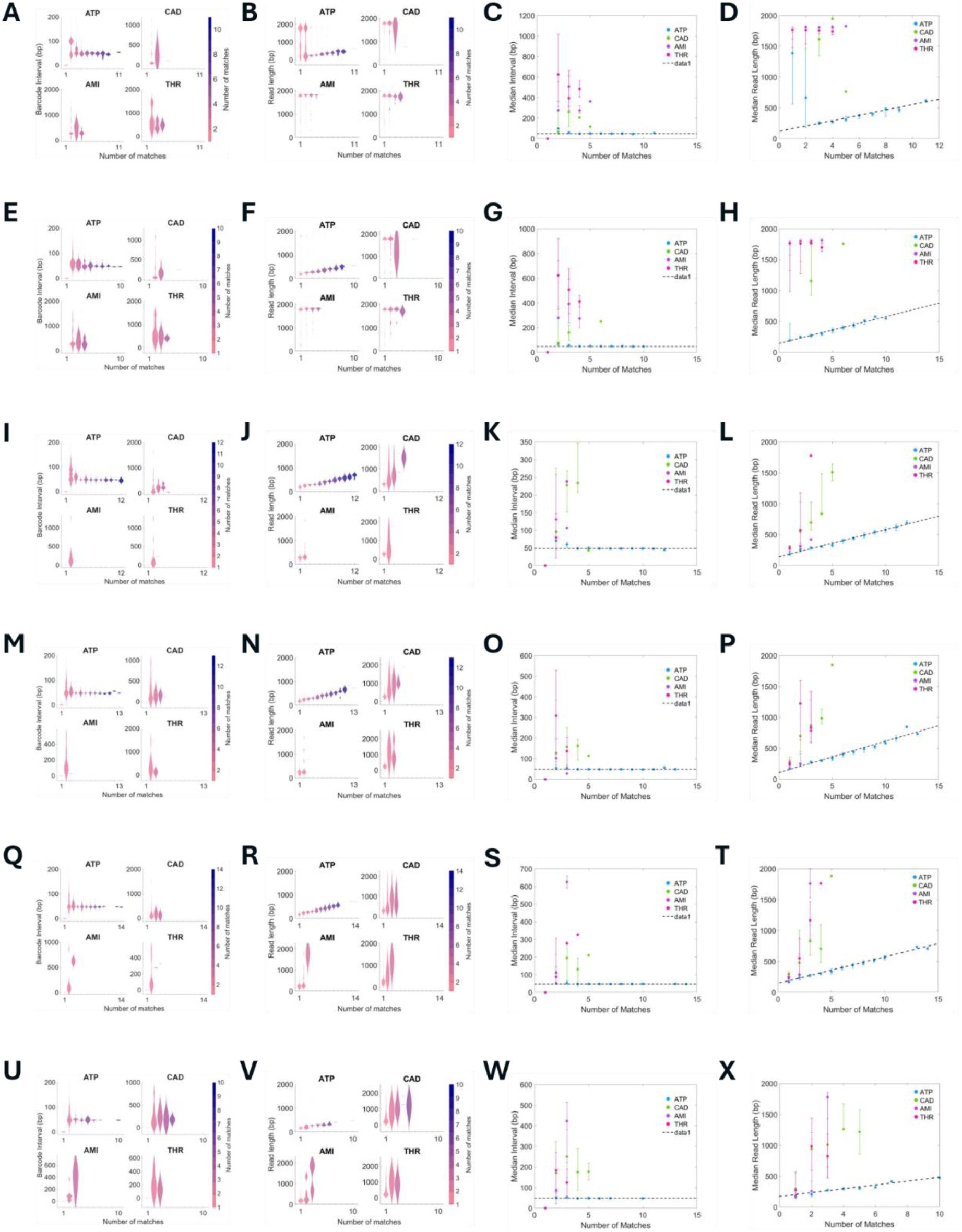
Barcode-interval and read-length distributions for the ATP singleplex replicates. (A, E, I, M, Q, U) Barcode interval (nucleotides) as a function of the number of matches for the specific target (ATP) and the non-specific targets (miR-AMI, miR-CAD, THR) in replicates E1-E6. (B, F, J, N, R, V) Read length as a function of the number of matches for all targets in replicates E1-E6. (C, G, K, O, S, W) Median barcode interval as a function of the number of matches in replicates E1-E6. (D, H, L, P, T, X) Median read length as a function of the number of matches in replicates E1-E6.

**Supplementary Fig. S3:**
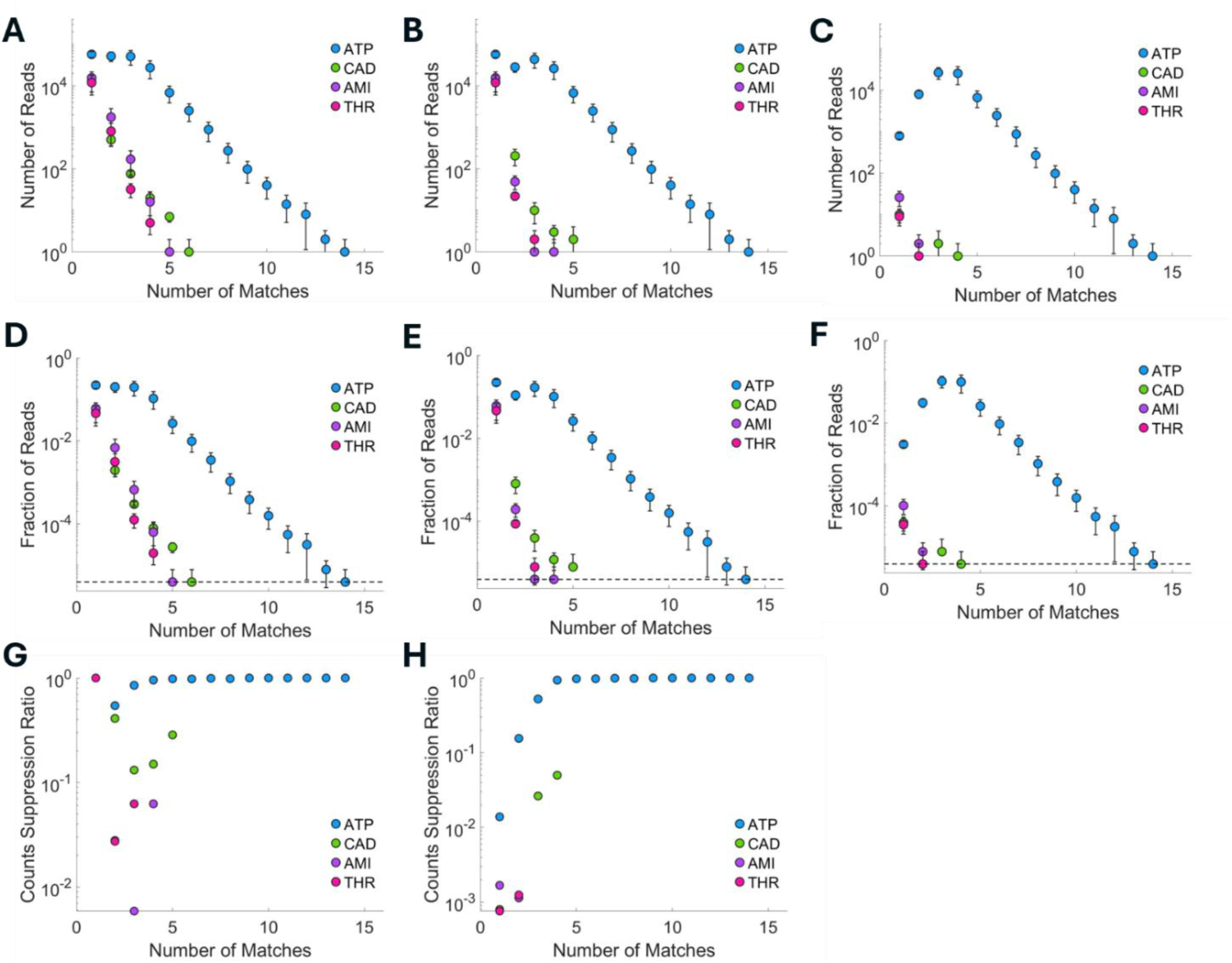
Filtered barcode-match statistics for the ATP singleplex assay. (A-C) Read count as a function of the number of matches: (A) without filtering, (B) after the periodicity filter, and (C) after the periodicity and read-length filters. (D-F) The corresponding fractional read counts: (D) without filtering, (E) after the periodicity filter, and (F) after both filters. (G, H) Suppression of read count achieved (G) by the periodicity filter alone and (H) by the periodicity and read-length filters together.

#### miR-AMI / miR-CAD multiplexed assay

**Supplementary Fig. S4:**
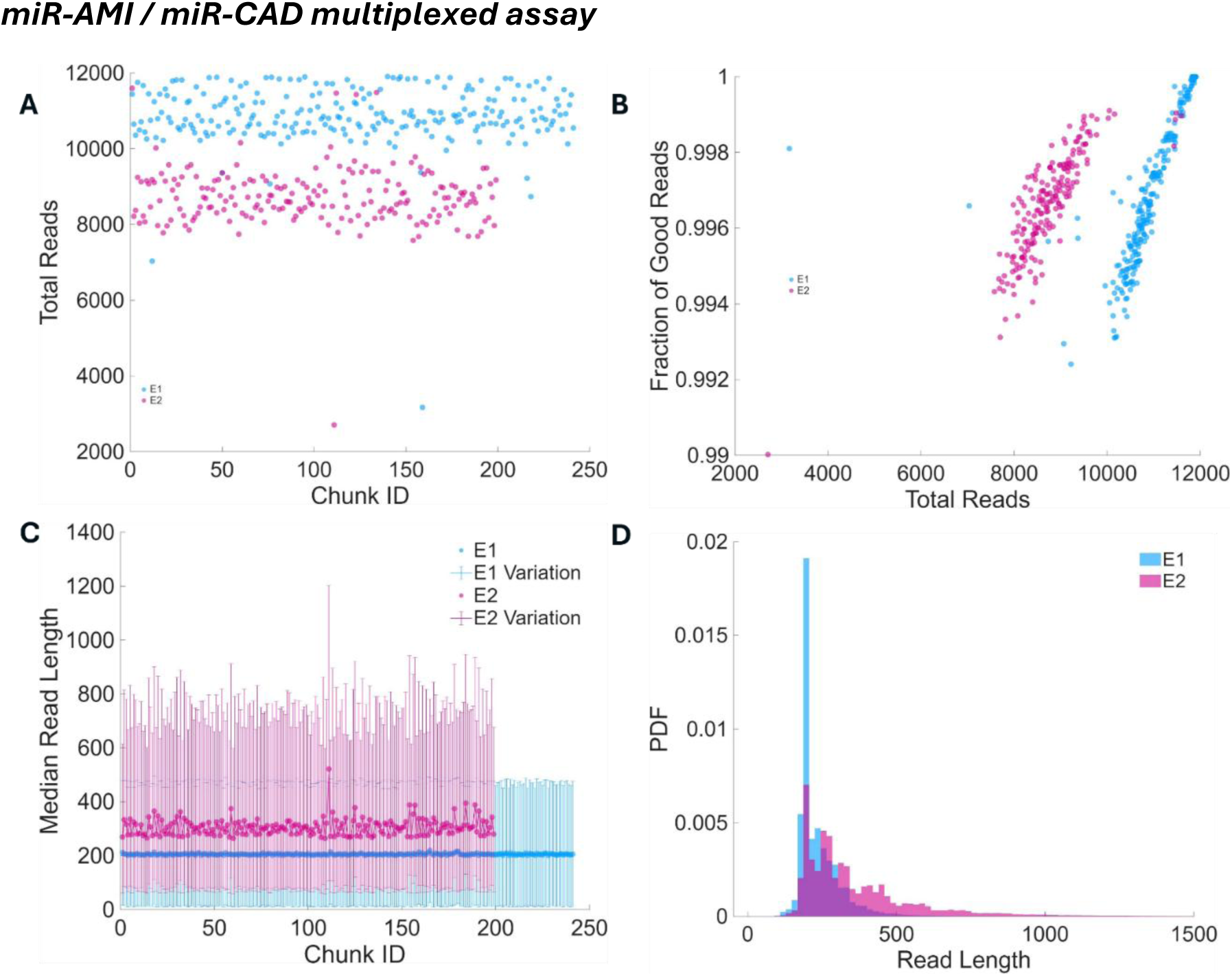
Read-length statistics for the miR-AMI/miR-CAD replicates. (A) Total reads per chunk for all chunks across both replicates. (B) Fraction of good reads for each replicate. (C) Median read length per chunk for both replicates. (D) Probability density of read length for both replicates.

**Supplementary Fig. S5:**
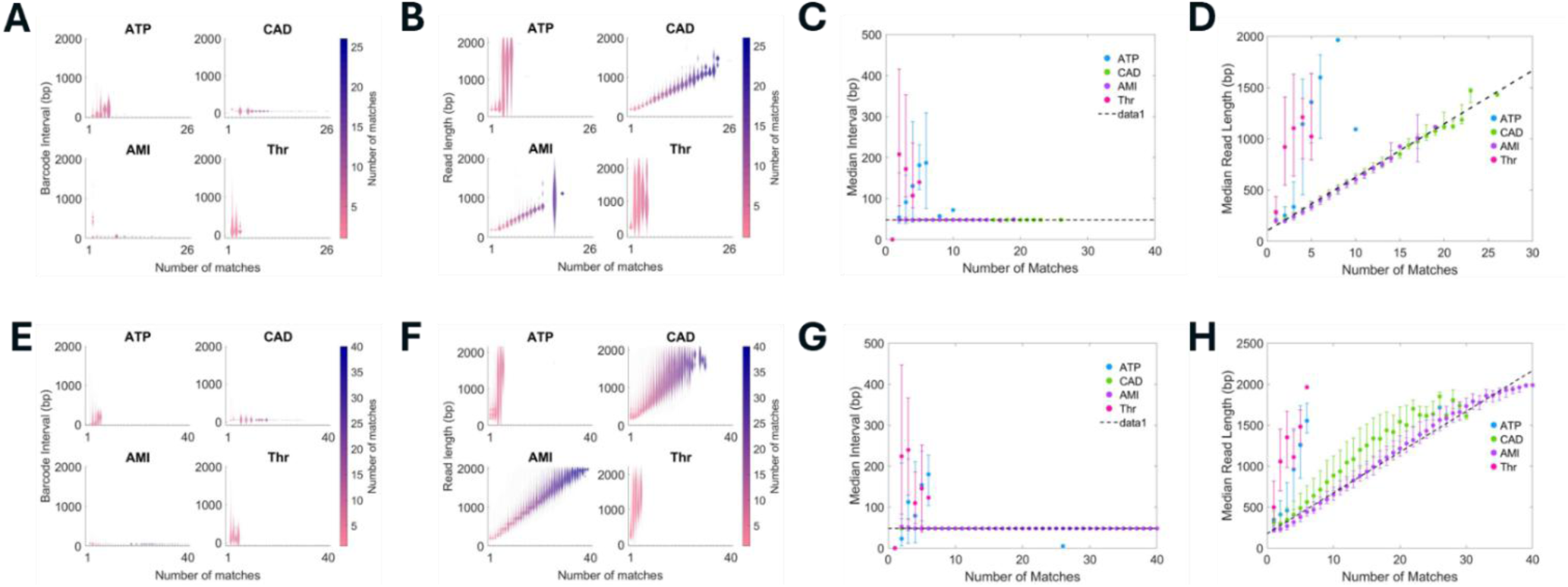
Barcode-interval and read-length distributions for the miR-AMI/miR-CAD replicates. (A, E) Barcode interval (nucleotides) as a function of the number of matches for the specific targets (miR-AMI, miR-CAD) and the non-specific targets (ATP, THR) in both replicates. (B, F) Read length as a function of the number of matches for all targets in both replicates. (C, G) Median barcode interval as a function of the number of matches in both replicates. (D, H) Median read length as a function of the number of matches in both replicates.

**Supplementary Fig. S6:**
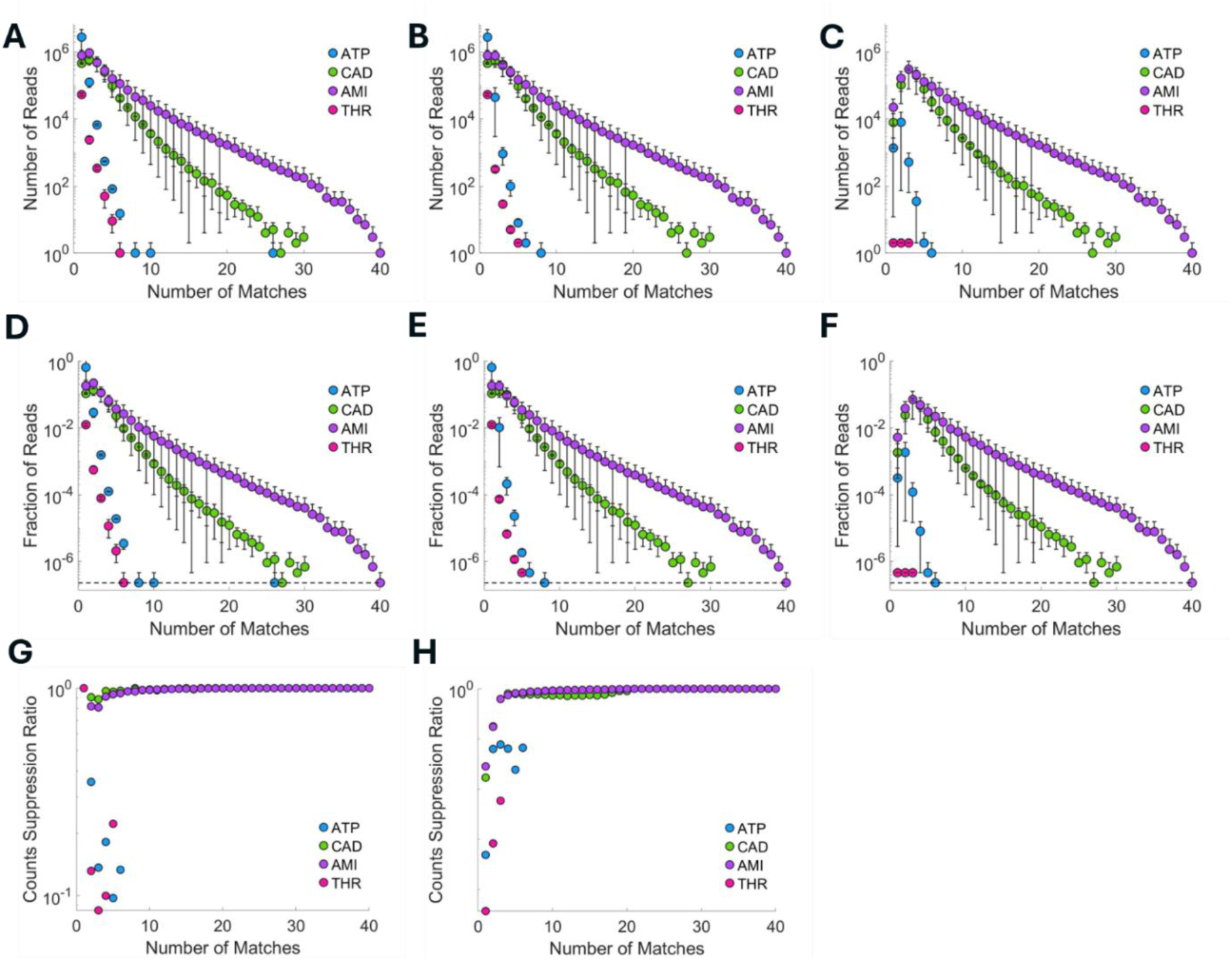
Filtered barcode-match statistics for the miR-AMI/miR-CAD assay. (A-C) Read count as a function of the number of matches: (A) without filtering, (B) after the periodicity filter, and (C) after the periodicity and read-length filters. (D-F) The corresponding fractional read counts. (G, H) Suppression of read count achieved (G) by the periodicity filter alone and (H) by both filters together.

**Supplementary Fig. S7:**
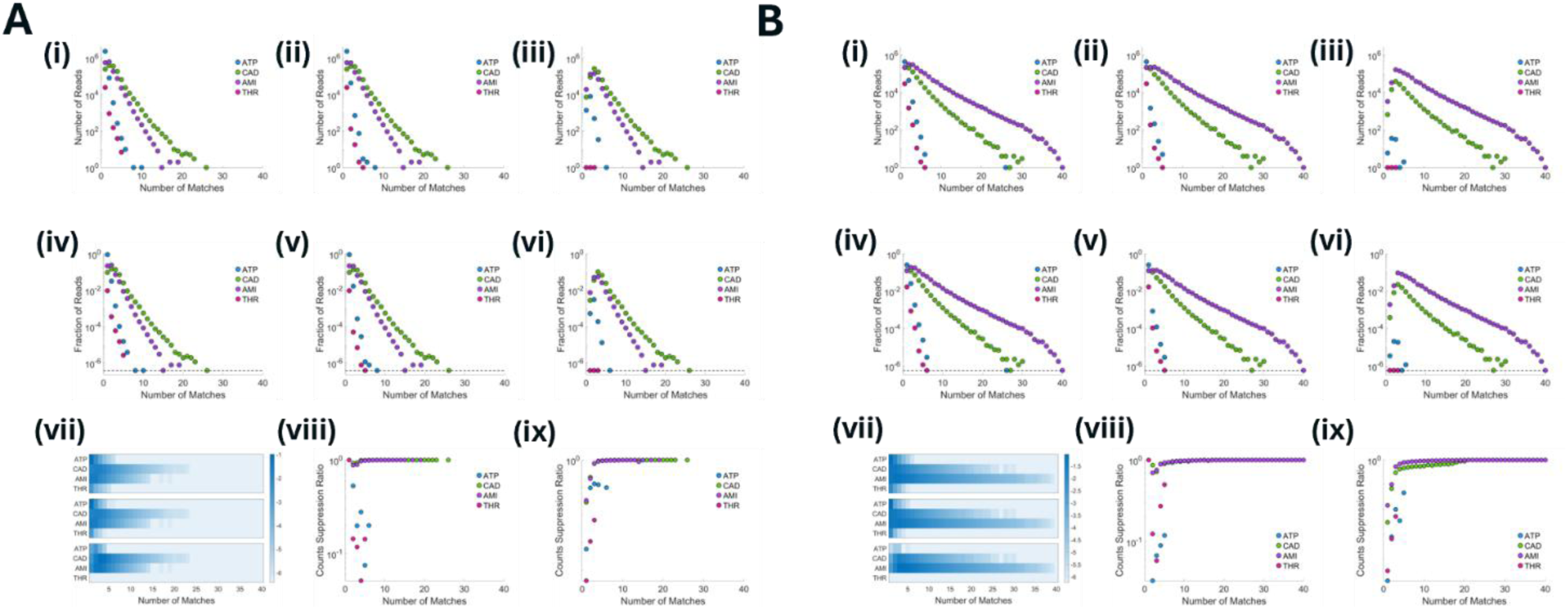
Per-replicate barcode-match statistics for the miR-AMI/miR-CAD assay. Panel (A) corresponds to the first replicate and panel (B) to the second. Within each panel: (i-iii) read count as a function of the number of matches without filtering, after the periodicity filter, and after both filters; (iv-vi) the corresponding fractional read counts; (vii) two-dimensional maps of the barcode-match statistics without filtering, after the periodicity filter, and after both filters; and (viii, ix) suppression of read count by the periodicity filter alone and by both filters together.

#### miR-AMI / miR-CAD / Thrombin multiplexed assay

**Supplementary Fig. S8:**
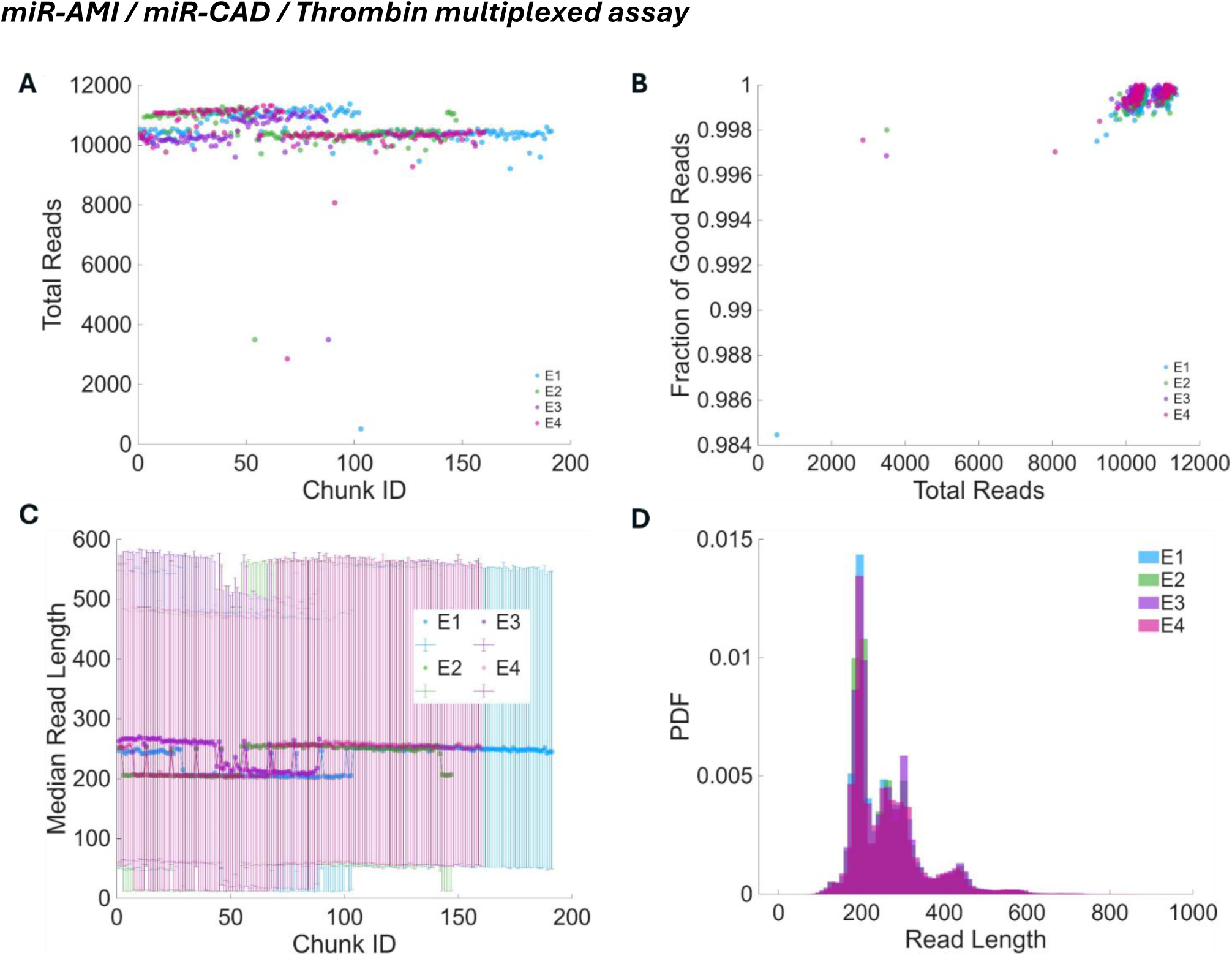
Read-length statistics for the miR-AMI/miR-CAD/THR replicates. The assay was performed in four replicates. (A) Total reads per chunk for all chunks across the four replicates. (B) Fraction of good reads for each replicate. (C) Median read length per chunk for all replicates. (D) Probability density of read length for the four replicates.

**Supplementary Fig. S9:**
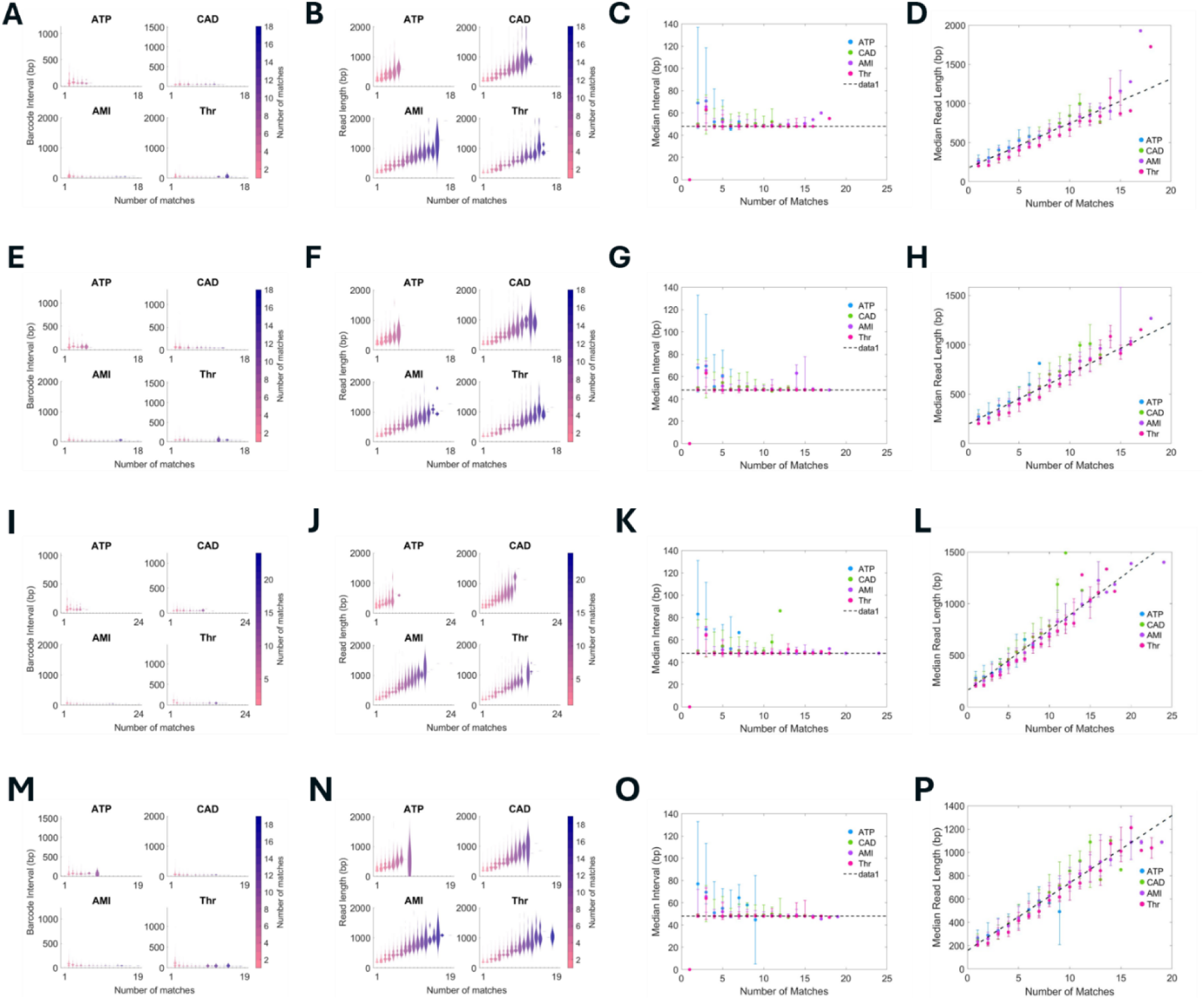
Barcode-interval and read-length distributions for the miR-AMI/miR-CAD/THR replicates. (A, E, I, M) Barcode interval (nucleotides) as a function of the number of matches for the specific targets (miR-AMI, miR-CAD, THR) and the non-specific target (ATP) in the four replicates. (B, F, J, N) Read length as a function of the number of matches for all targets in the four replicates. (C, G, K, O) Median barcode interval as a function of the number of matches in the four replicates. (D, H, L, P) Median read length as a function of the number of matches in the four replicates.

**Supplementary Fig. S10:**
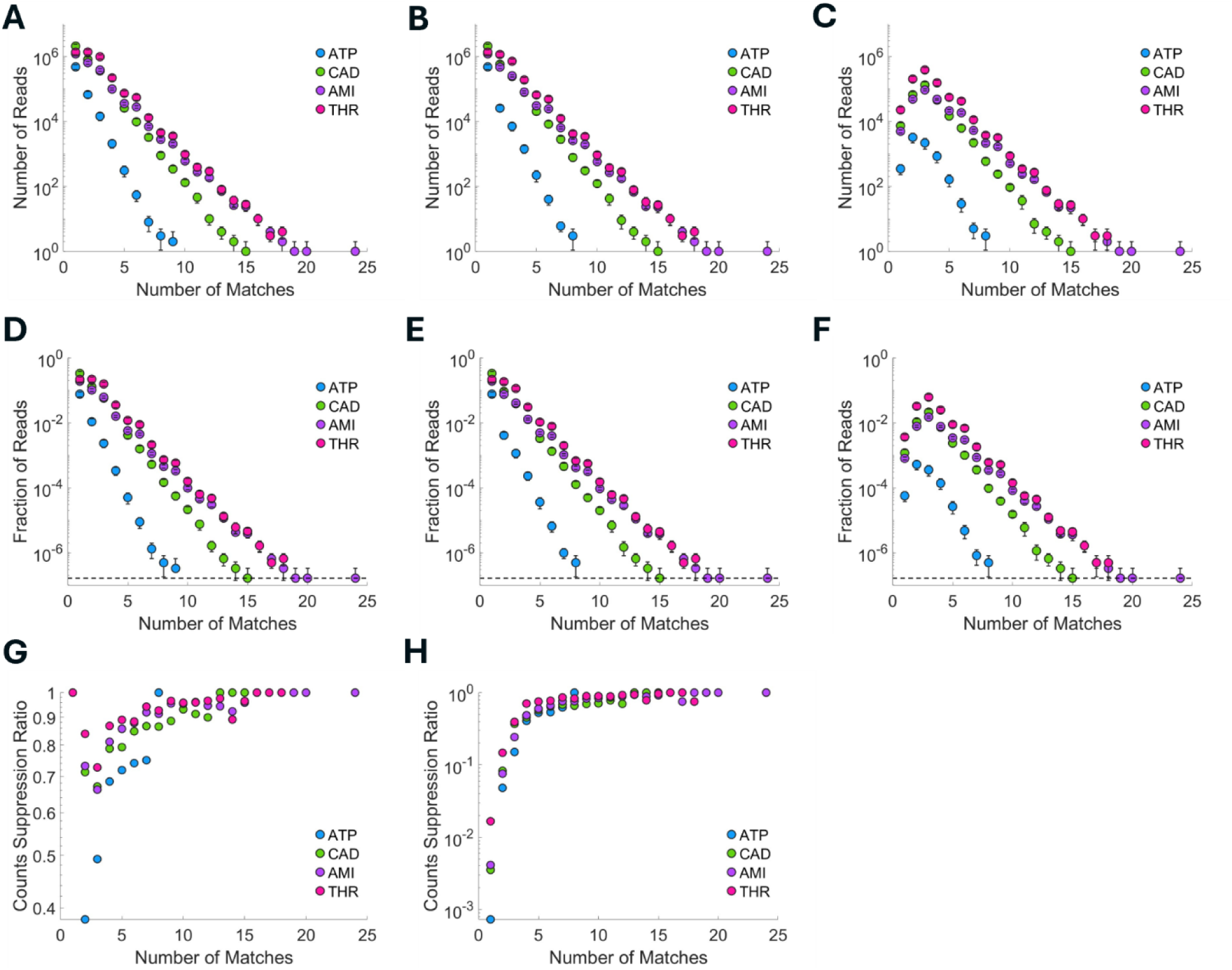
Filtered barcode-match statistics for the miR-AMI/miR-CAD/THR assay. (A-C) Read count as a function of the number of matches: (A) without filtering, (B) after the periodicity filter, and (C) after the periodicity and read-length filters. (D-F) The corresponding fractional read counts. (G, H) Suppression of read count achieved (G) by the periodicity filter alone and (H) by both filters together.

**Supplementary Fig. S11:**
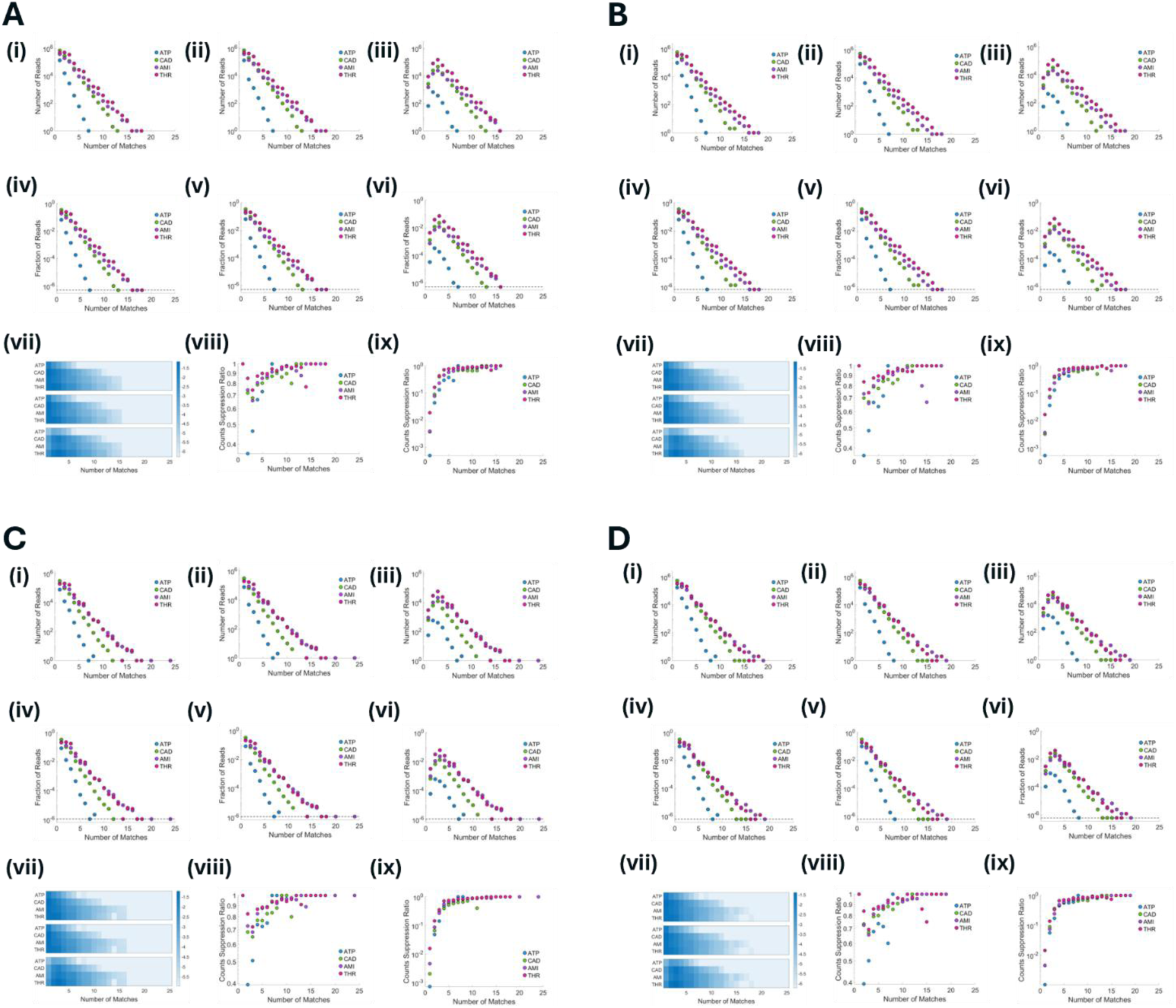
Per-replicate barcode-match statistics for the miR-AMI/miR-CAD/THR assay. Panels (A-D) correspond to the four replicates. Within each panel: (i-iii) read count as a function of the number of matches without filtering, after the periodicity filter, and after both filters; (iv-vi) the corresponding fractional read counts; (vii) two-dimensional maps of the barcode-match statistics without filtering, after the periodicity filter, and after both filters; and (viii, ix) suppression of read count by the periodicity filter alone and by both filters together.

#### MinION miR-CAD singleplex assay

**Supplementary Fig. S12:**
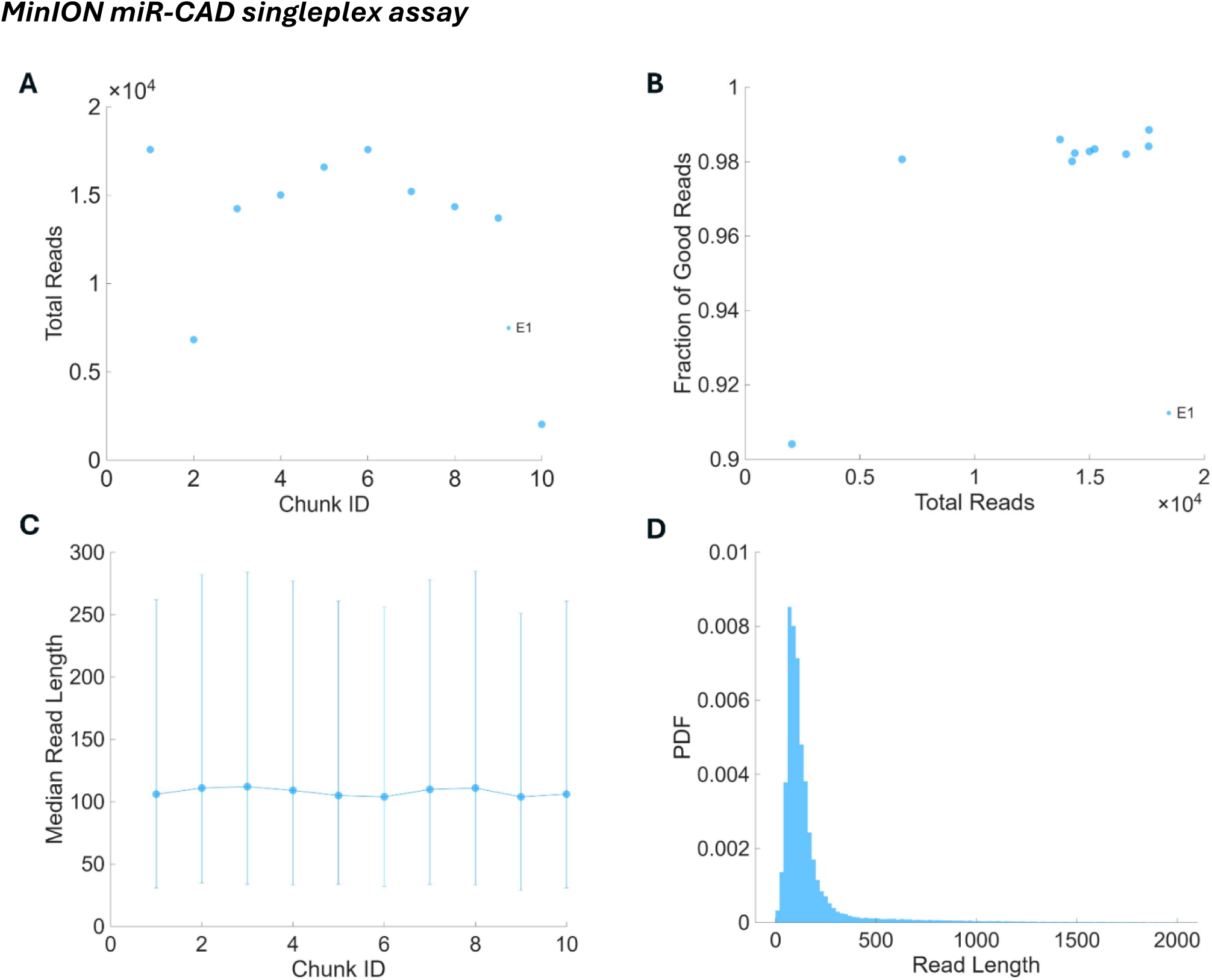
Read-length statistics for the MinION miR-CAD singleplex assay. (A) Total reads per chunk. (B) Fraction of good reads. (C) Median read length per chunk. (D) Probability density of read length.

**Supplementary Fig. S13:**
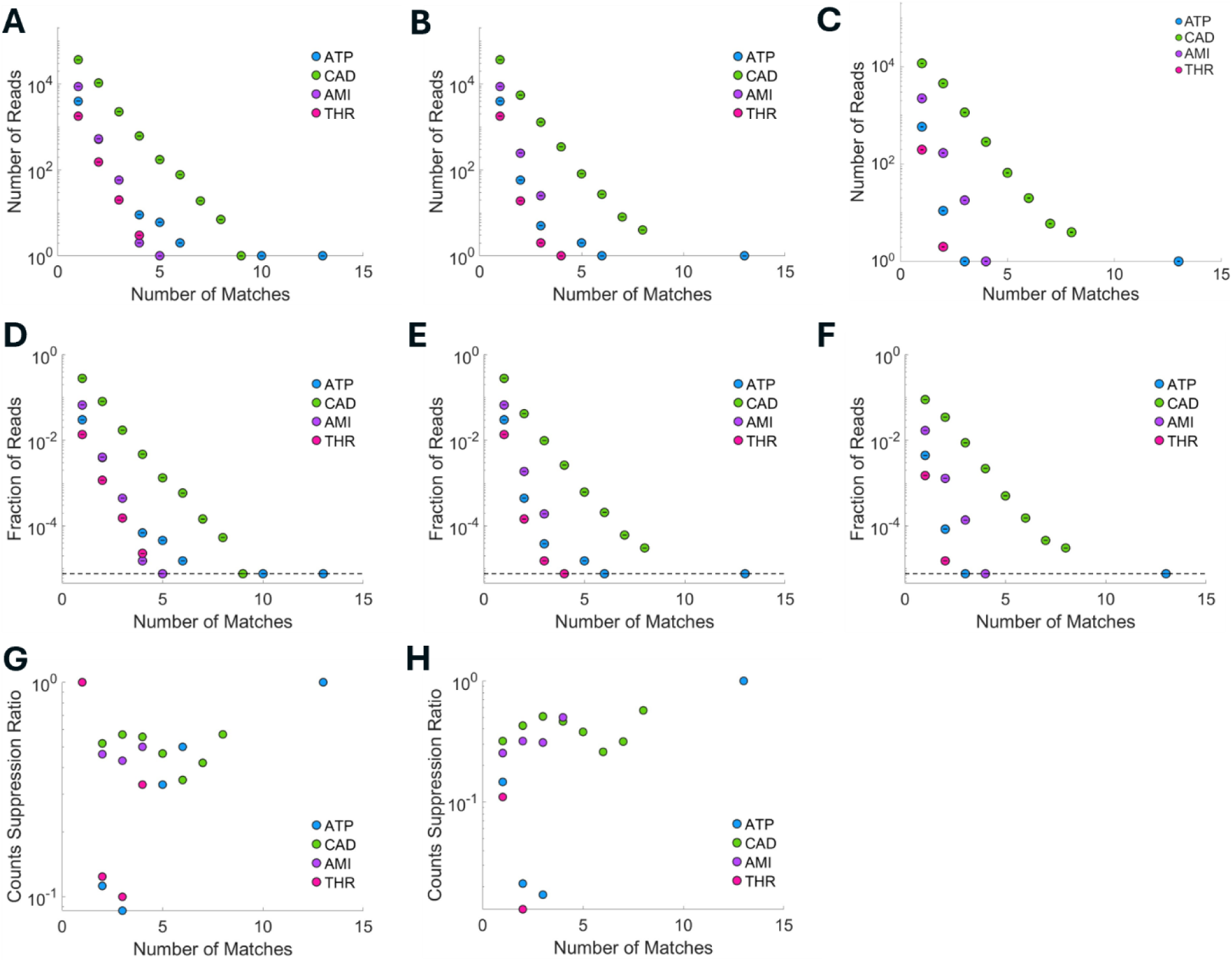
Filtered barcode-match statistics for the MinION miR-CAD singleplex assay. (A-C) Read count as a function of the number of matches: (A) without filtering, (B) after the periodicity filter, and (C) after the periodicity and read-length filters. (D-F) The corresponding fractional read counts. (G, H) Suppression of read count achieved (G) by the periodicity filter alone and (H) by both filters together.

#### Detection in a bacterial genome background (E. Coli K-12)

**Supplementary Fig. S14:**
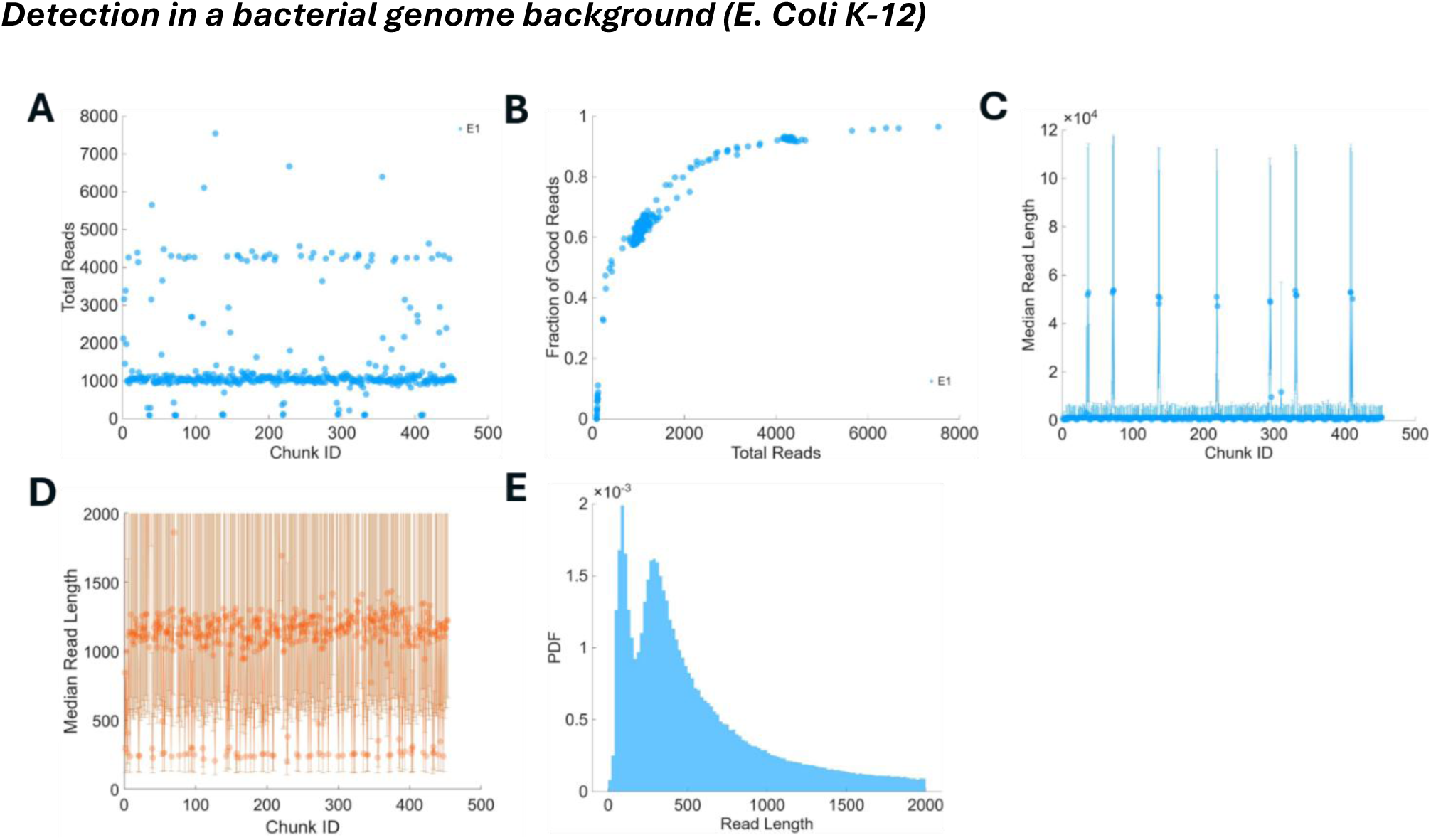
Read-length statistics for miR-AMI detection in the bacterial whole-genome background. (A) Total reads per chunk. (B) Fraction of good reads. (C) Median read length per chunk. (D) Median read length per chunk, shown on an expanded scale. (E) Probability density of read length.

**Supplementary Fig. S15:**
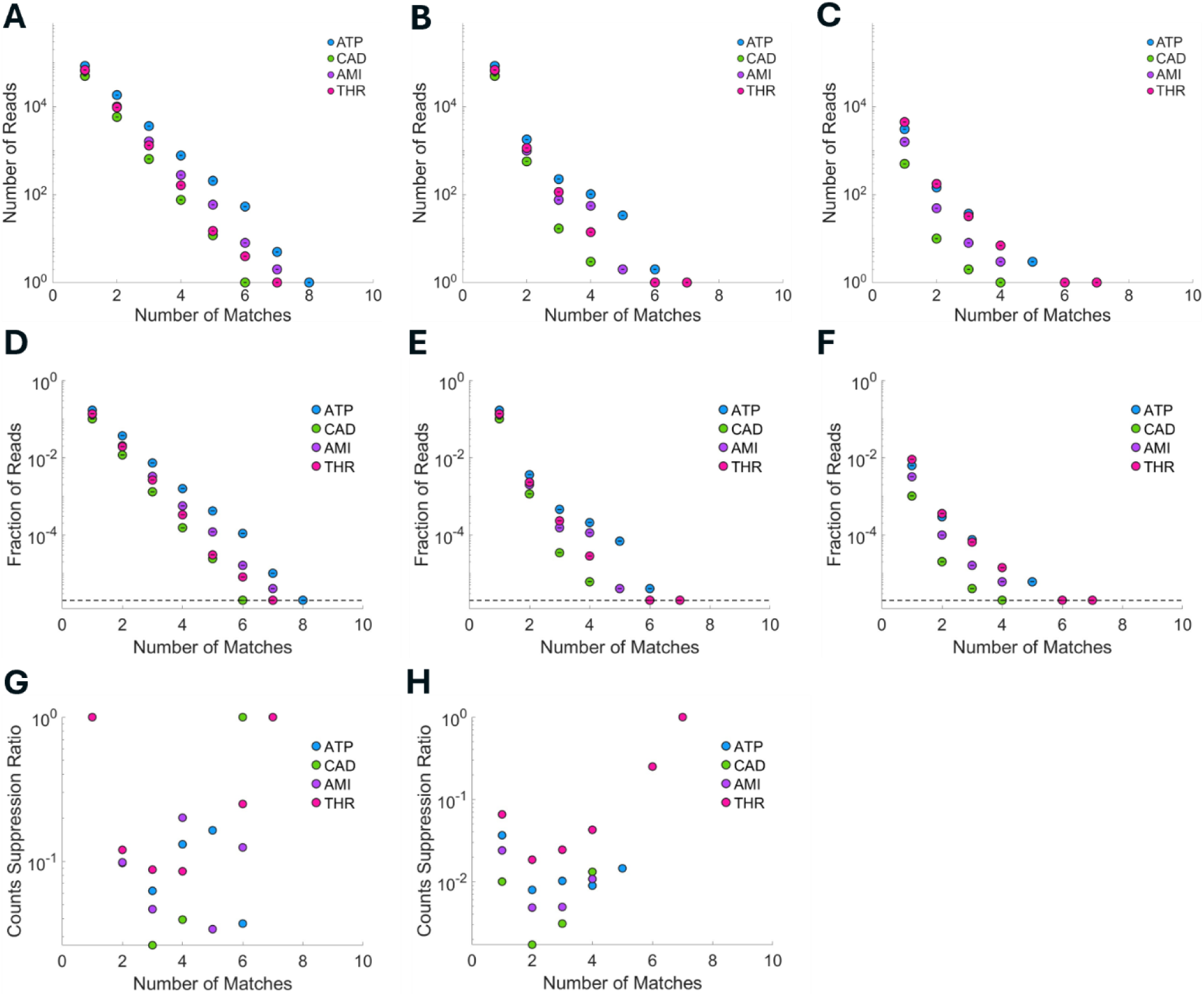
Filtered barcode-match statistics for miR-AMI detection in the bacterial whole-genome background. (A-C) Read count as a function of the number of matches: (A) without filtering, (B) after the periodicity filter, and (C) after the periodicity and read-length filters. (D-F) The corresponding fractional read counts. (G, H) Suppression of read count achieved (G) by the periodicity filter alone and (H) by both filters together. Residual signal is observed for the non-specific targets, indicating that the 10-bp barcode is insufficient in this matrix.

### 3. Materials and Methodology

All the DNA sequences were procured at 100µM stock, as HPLC-purified single-stranded DNA oligonucleotides from Integrated DNA Technologies (IDT), which were then folded to form hairpins. DNA sequences were designed with possible secondary structures in mind. The reaction buffer used for Hybridization Chain Reaction (HCR) and for formation of hairpins contained 5mM MgCl2/ 0.3 M NaCl/ 20 mM Tris (pH 7.6). For folding, 10µM single-stranded DNA oligonucleotide resuspended in the reaction buffer was incubated at 95⁰C for 1 minute, followed by quick cooling on ice for 90 minutes.

For all the assays described in the manuscript, 3µL each of 10µM initiator, H1, and H2 hairpins form each hairpin sets for ATP, miR-CAD, miR-AMI, and Thrombin are present in the reaction mixture. For the ATP singleplex assay, the hairpin sets are added to 20µL of reaction buffer, making the total reaction volume 56µL. To this, 3µL of 16.5 mM ATP stock solution (from ATP Bioluminescence Assay Kit CLS II from Roche) was added to make a final ATP concentration of approximately 1mM. For the miR-AMI/miR-CAD multiplexed detection, the hairpin sets for the four targets were added to 110µL of the reaction buffer. To this, 3µL each of 10µM miR-AMI and miR-CAD was added, to make a final concentration of approximately 200nM of miR-AMI and mi-CAD. For the miR-AMI/miR-CAD/THR triplex, the hairpin sets for the four targets were added to 110 µL of the Reaction buffer. To this, 3µL each of 10µM miR-AMI, miR-CAD and 50U/µL Thrombin was added, to make a final concentration of approximately 200nM for miR-AMI and miR-CAD, and approximately 1U/µL for Thrombin. For the cross-platform validation through single-plex detection of miR-CAD, the hairpin sets for the four targets were added to 75µL of the Reaction buffer. To this, 8µL of 10µM miR-CAD was added, to make a final concentration of approximately 670nM for miR-CAD. The *Escherichia coli* str. K-12 substr. MG1655 (GenBank Accession number: CP174267.1) genomic DNA was isolated using a commercial kit as per the vendor protocol [R1]. The hairpin sets for the four targets were added to 100µL of 80ng/µL of extracted genomic DNA. To this, 3µL of 10µM miR-AMI was added, to make a final concentration of approximately 200nM for miR-AMI.

All assay reactions were incubated at Room temperature for 24 hours, before running on agarose gel to qualify for downstream Long-Read Sequencing (LRS) on the Oxford Nanopore platform. The DNA concatemer library formed after HCR was library prepared with Oxford Nanopore Technologies Ligation Sequencing DNA V14 (SQK-LSK114) protocol, involving DNA repair and end-repair, followed by Adapter ligation [R2]. The Promethion sequencing was done at the external site [R3]. For in-house sequencing, 75µL of the DNA library was loaded onto the MinION flow cells. Base-calling was done using Dorado with Super High Accuracy (SUP) model [R4].

Gel electrophoresis was used as a preliminary characterization method to verify the success of HCR amplification.1% agarose gel containing 0.5µg/mL EtBr was prepared in 1X SB buffer (10mM NaOH, pH adjusted to 8.5 with Boric acid). Electrophoresis was done at 100V for 1.5 hours in 1X SB running buffer. Figure S16 shows miAMI (Panel A) and miCAD (Panel B) HCR reaction products at different concentrations including a negative control, i.e. without the target. In these reactions, the HCR set corresponding to the specific target was added along with the target at different concentrations. One can observe discrete bands indicative of successful HCR reaction in the presence of the target (Lanes 3, 4, and 5 in both panels) whereas the negative controls (Lane 2 in both panels) do not show strong discrete bands that are produced due to HCR. Further comparison with the 1 kb DNA ladder (NEB) shows the majority of the HCR products lies below 2 kbp in length, justifying the length filter mentioned in the main text.

**Supplementary Fig. S16:**
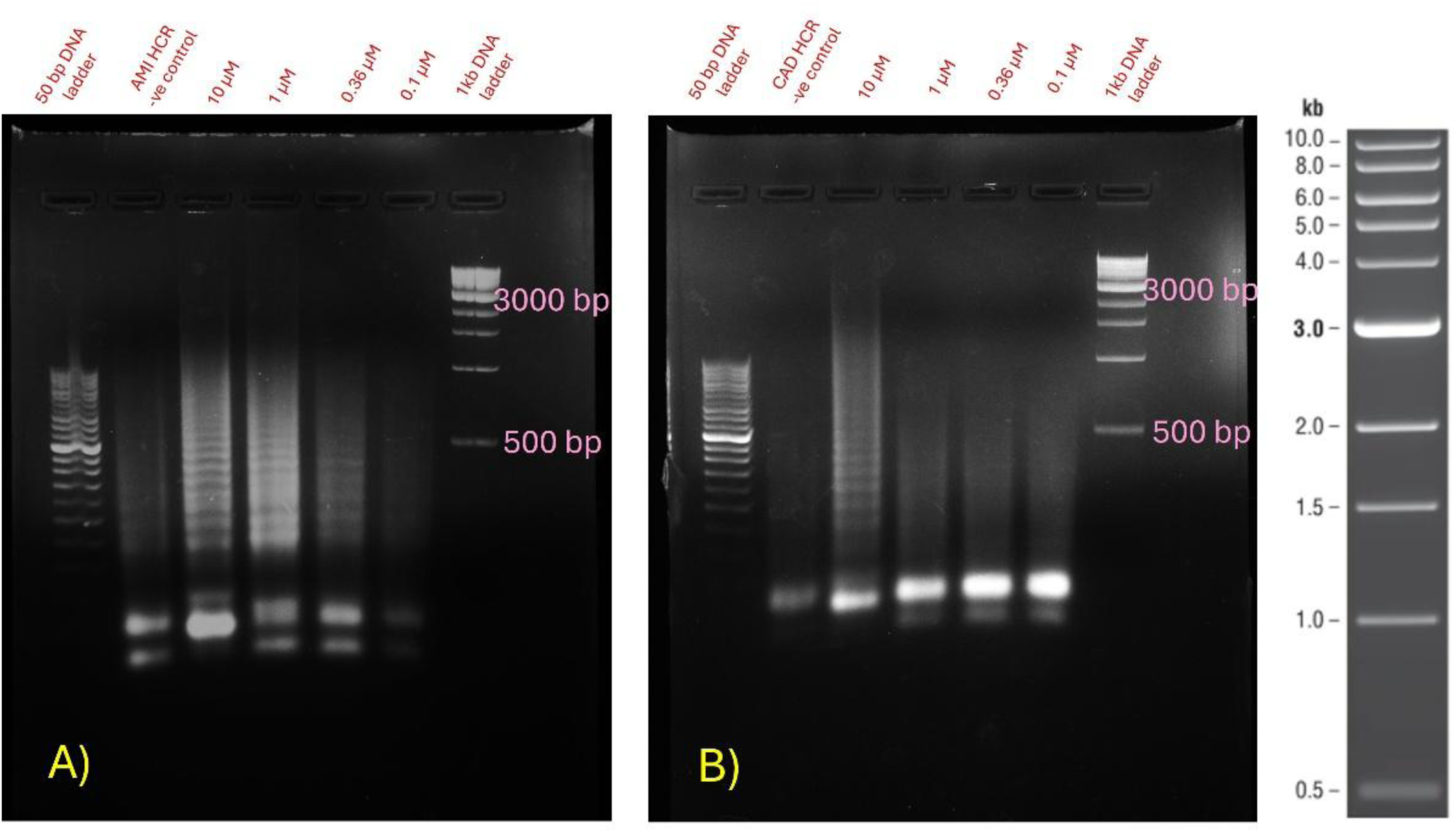
Gel-electrophoresis of HCR products (A) Gel image of HCR reaction with miAMI as the target at 4 different concentrations. The presence of the target, miAMI results in discrete bands (Lane 3, 4 and 5) that result from incorporation of periodic blocks as shown in the schematic Figure 1 in the main text. The bands are in the range of 50 – 2000 bp by comparison with the 1kb DNA ladder used in the experiment. (B) Gel image of the HCR reaction with miCAD as the target at 4 different concentrations indicating identical behavior.

### 4. Flanking-Sequence Analysis of Periodic Barcodes in HCR Sequencing Reads

Sequencing reads generated by the aptamer-triggered hybridization chain reaction (HCR) assay were analyzed to characterize the sequence context immediately flanking the target-encoding barcode. Each of the four targets was assigned a unique ten-nucleotide DNA barcode (SI Table 1). As the HCR product is a periodic concatemer, the barcode will recur at a characteristic spacing along each read. For each target, barcodes were identified as described in the main text (Decoding Framework section) with further details provide below.

Reads were selected according to two criteria. First, the measured mean barcode interval *λ_read_* should lie within a window *w_λ_* of the designed period, *λ_des_* = 48 *nt* i.e., |*λ_read_* − *λ_des_*| < *w_λ_*, thereby retaining only those reads in which the barcode recurred with the spacing expected of the HCR product. Second, a length-consistency criterion, (*L_read_*(*m*) − *L*_0_ − *λ_des_m_read_*) < *w_L_*, was imposed, in which *L_read_* denotes the read length, *m_read_* is the number of barcode matches detected, and *L*_0_ is the constant linear fitting term that represents the pre-barcode length fixed by the design. This condition requires the matches to tile the read densely and suppresses reads in which sparse, isolated matches would otherwise satisfy the interval filter. Reads meeting both criteria were retained. Within each retained read, barcode occurrences were localised by an exhaustive, gapless sliding-window scan. A window of width equal to the barcode length (10 nt) was advanced one nucleotide at a time across the read, and at each position the number of mismatches between the window and the barcode was determined. Positions at which the barcode matched within a Hamming distance of two, with no insertions or deletions permitted, were recorded as barcode occurrences. Reads yielding number of occurrences more than expected from the read-length cutoff of 2,000 were discarded as probable artefacts. As the maximum number of occurrences expected is 2000/48 ∼ 40, 40 was used as an upper limit for the number of occurrences.

For each recorded occurrence, the flanking sequences, specifically, six nucleotides immediately upstream of the barcode (the prefix) and eight nucleotides immediately downstream of the ten-nucleotide barcode region (the suffix), were extracted. Positions extending beyond either terminus of the read were padded with a gap character to preserve register. The extracted flanks were indexed by the ordinal position of the barcode occurrence within the read, so that the context of the first, second, and subsequent barcode repeats could be aggregated separately, and were pooled across all reads. For a given ordinal position, the pooled prefix and suffix sequences were each sorted independently in alphabetical order to group similar motifs. The sorted collection was then rendered as a nucleotide-resolved alignment image, in which each sequence occupies one row, each column corresponds to a position within the flank, and the four nucleotides and the gap character are distinguished by color. This representation clearly visualizes the conserved and variable positions of the flanking sequences adjacent to the barcode.

Figure S17 below shows the flanking sequences for number of matches, k = 1 and k = 25, for the AMI barcodes in the AMI-CAD two-plex assay (replicate E2). The most dominant suffix sequence for both k = 1 and k = 25, is TCTTGTTA. From SI Table 2, we see that this sequence is identical to that of Block *B*_0_ (SI Table 2, Figure 1 in main text). However, there appears to be two dominant prefix sequences for both k = 1 and k = 25, occurring at relatively similar abundance. They are, namely, GAGATG and CATCTC. Again, referring to SI Table 2, we see that these correspond to the block *A* and 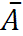 (Figure 1). Referring to Figure 1 in the main text, we can see that the 8 nt long suffix sequence of the barcode region will be *B*_0_ irrespective of whether the top or the bottom strand is read. However, the 6 nt long prefix sequence will correspond to the block *A* when the top strand is read and to block 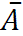 when the bottom strand is read. The number of reads with both of these blocks is roughly equal indicating that there is no significant bias in whether the top or the bottom strand is read. Here top and bottom strands refer to the schematic shown in Figure 1 of the main text. The results for ATP in the ATP single-plex (Figure S18) and for THR in the AMI-CAD-THR 3-plex (Figure S19) indicate the same conclusion across multiple targets, and number of occurrences, even with low number of reads (Figure S18C and D). The rest of the targets in all the assays also show the same behavior and are not shown here separately due to the large number of images required and also because they all point to the same conclusion.

**Supplementary Figure S17:**
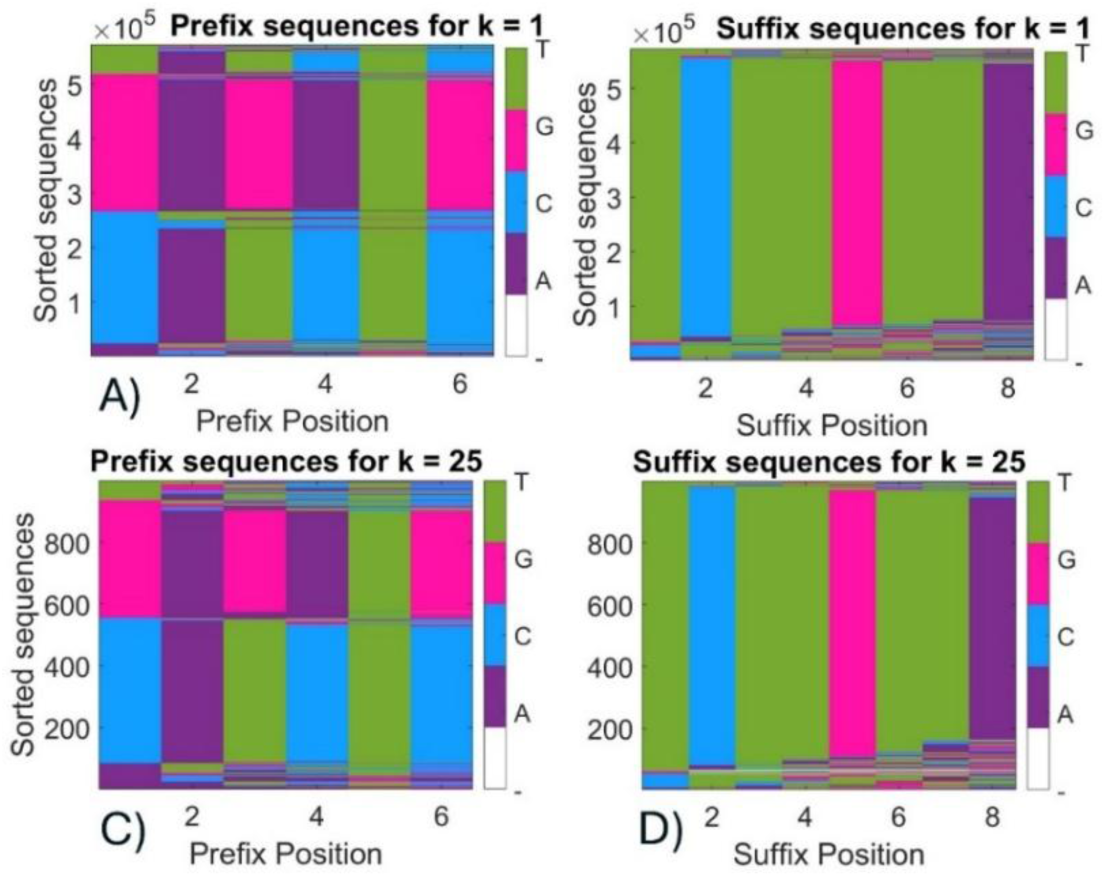
Flanking regions around the barcode for AMI in the AMI-CAD two-plex assay. Prefix (6 nt before the barcode start) and Suffix (8 nt after the barcode end) show that there is no bias in the sequencing reads between the top and bottom strand of the HCR product.

**Supplementary Figure S18:**
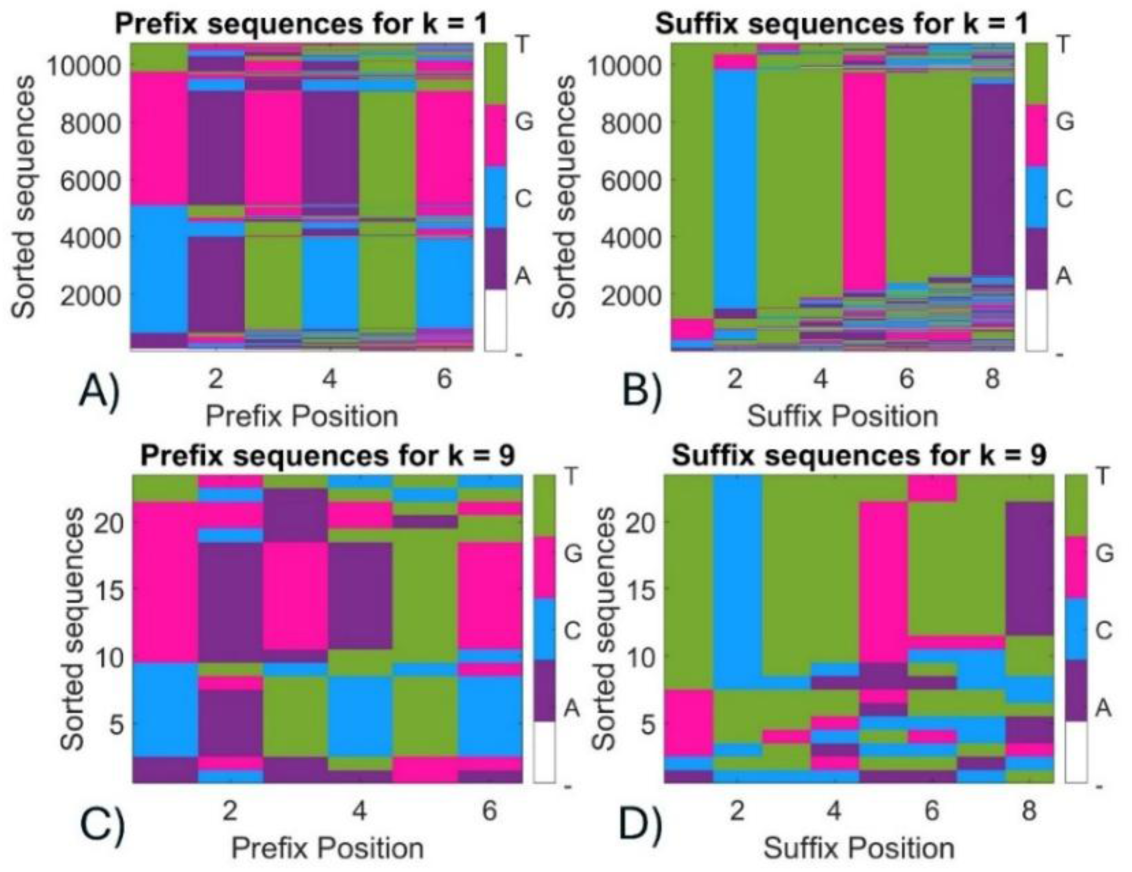
Flanking regions around the barcode for ATP in the ATP single-plex assay.

**Supplementary Figure S19:**
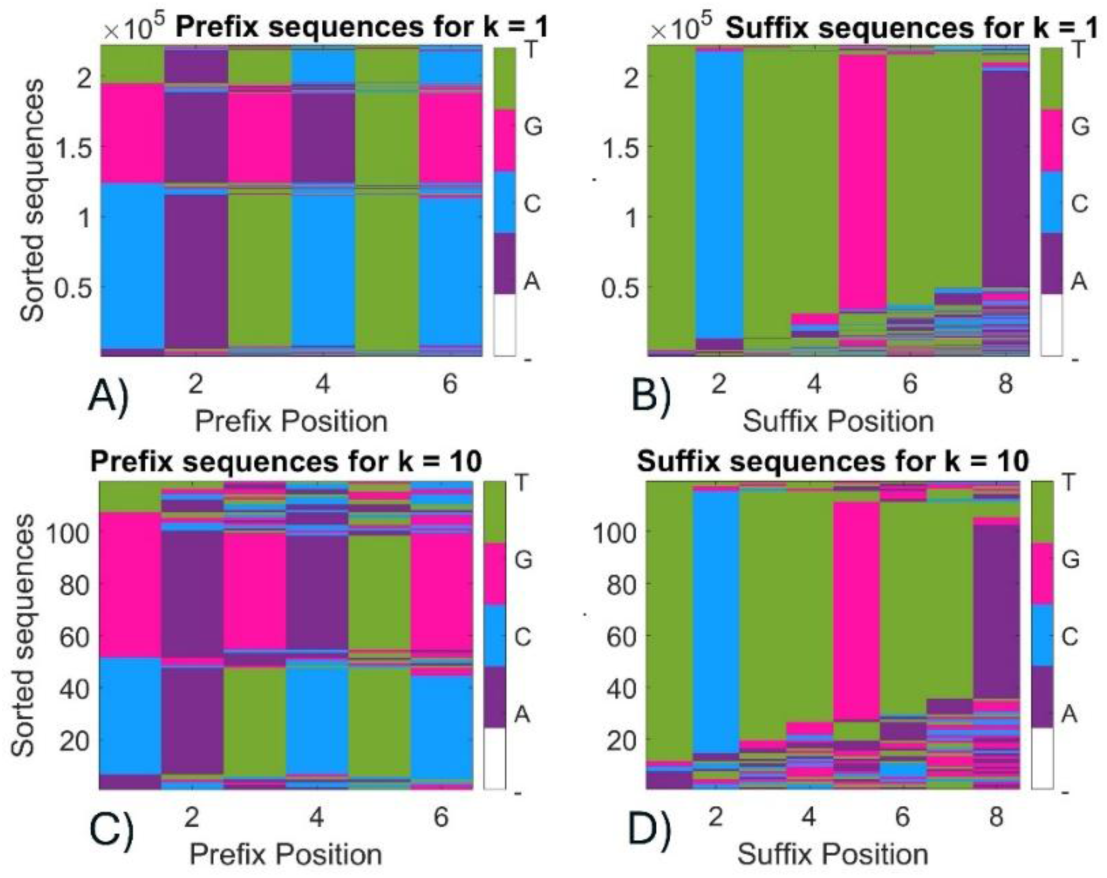
Flanking regions around the barcode for THR in the AMI-CAD-THR three-plex assay.

### 5. Stochastic Simulation and Mean-Field Theory of Target-Activated Initiator Barcoding of HCR Products

5.1 Notation used for the main variable
| Symbol | Quantity/Parameter |
| --- | --- |
| $I_0, T_0, B_0$ | Initial numbers of initiator, target, and barcode molecules, respectively |
| $I(t), I^*(t)$ | Numbers of inactive and activated initiators, respectively, at time $t$ |
| $T(t), B(t)$ | Numbers of target and barcode molecules at time $t$, respectively |
| $\kappa_a, \kappa_b$ | Activation and barcode-attachment rate constants, respectively |
| $N_a, N_b$ | Mean activation and barcode attachment events per step, respectively |
| $\alpha, \beta$ | Dimensionless activation-rate factor (activfrac) and barcoding-rate factor (bcratefrac), respectively |
| $\tau$ | Target fraction (tfrac), with $T_0 = \tau I_0$ |
| $s_j$ | Integer state of initiator $j$ |
| $n_{a,i}, n_{b,i}$ | Number of activation and barcode attachment events, respectively, in step $i$ |
| $\lambda$ | Mean number of barcodes per activated initiator |
| $N(k)$ | Expected number of initiators bearing exactly $k$ barcodes |
| $k_{max}$ | maximum tracked barcode count |
| $S_{steps}$ | Number of simulation steps |

#### 5.1 Notation used for the main variable

#### 5.2 Mean-field model of the barcoding reaction scheme

We created a reduced model of the multi-step reaction scheme shown in Figure 1 of the main text. The two salient features of this reaction scheme is the unbinding of the initiator hairpin (hairpin I in Main text Figure 1) due to the binding of the target. This allows the other two hairpins (H_1_ and H_2_ in the same figure) to undertake the cascade reaction leading to the formation of the concatemer with periodic insertions of the barcode. We made a reduced representation this reaction cascade using two reactions. The first one represents the “activation” of the initiator due to binding of the target molecule. The second reaction represents the binding or incorporation of barcodes to the activated initiators.

The activation reaction is represented by

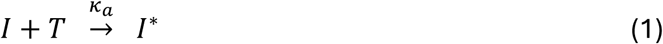

Here, *I* and *T* are the concentrations of the (un-activated) initiator molecules and target molecules, respectively; *I*^∗^ is the concentration of the activated initiators that are “activated” for barcode incorporation; and *κ_a_* is the reaction rate.

The barcode incorporation reaction is represented by

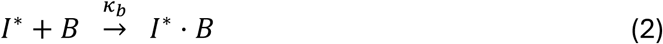

where *I*^∗^ denotes the concentration of the activated initiator, *B* is the concentration of barcodes available; *I*^∗^ · *B* is the initiator-barcode complex (concatemer in Figure 1 of the main text) and *κ_b_* is the barcode-attachment rate constant. The initial (time *t* = 0) concentrations the initiator, target and barcode are *I*_0_, *T*_0_, and *B*_0_, respectively and *I*^∗^(*t* = 0) = 0.

The time evolution of the concentration *I*(*t*), based on Eq. (1) and (2), will be governed by

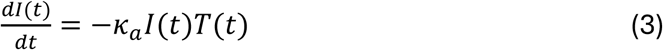

Conservation of initiators implies that

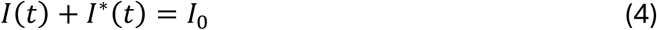

On similar grounds, conservation of target molecules implies

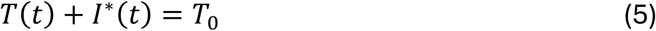

Using Eqs. (3), (4) and (5) reduces the activation kinetics to a single Riccati-type equation,

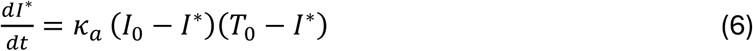

For *T*_0_ ≠ *I*_0_, defining *E*(*t*) = exp[*κ_a_*(*T*_0_ − *I*_0_) *t*], the solution subject to *I*^∗^(0) = 0 is

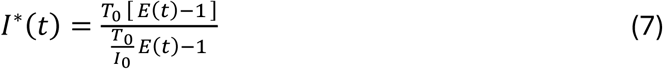

In the degenerate case *T*_0_ = *I*_0_, Eq.(7) reduces to

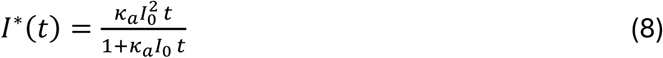

The barcode population follows from *dB*/*dt* = −*κ_b_B I*^∗^, giving

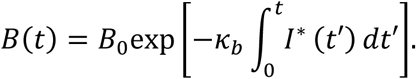

Substituting *I*^∗^(*t*) from Eq. (7), the required integral evaluates in closed form,

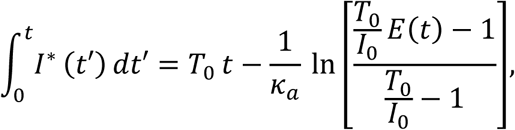

so that, for *T*_0_ ≠ *I*_0_,

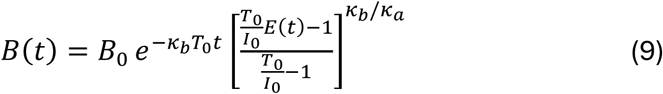

and, for *T*_0_ = *I*_0_, we use *I*^∗^(*t*) from Eq. (8), resulting in

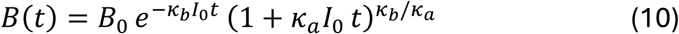

The model presented here is a continuum, mean-field model, while the experimental data is available in terms of discrete number of sequencing reads with *m* barcode matches. Therefore, a discrete stochastic model is required to map the theoretical model to experimental data. The primary value of the mean-field model presented here is to provide a deterministic reference for the stochastic trajectories of the species calculated as described in the next section.

#### 5.3 Stochastic simulation of the barcoding reaction scheme

In the stochastic regime, the concentrations of the species is replaced by number of copies, and time is measured in simulation steps of *i* = 0, 1, …, *S_steps_*.

Conservation of the number of initiators and targets demands that, at any time-step *i*

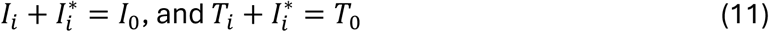

The activation step consumes one target per event, whereas barcode attachment increases the barcode count of an activated initiator without altering its activation status.

The mean number of activation events *N_a_* and the mean number of barcode attachment events *N_b_* are written in terms of two rate parameters *α* and *β* as follows

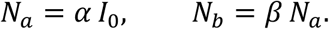

The rate constants for activation, *κ_a_* and barcode-attachment, *κ_b_* are then set as

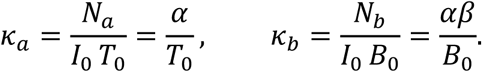

Consequently, *α* may be interpreted as the fraction of the initiator population activated during the first step at the initial target abundance.

The simulation scheme is as follows. Each initiator *j* is assigned an integer state *s_j_*. The convention adopted is *s_j_* = 0 for an inactive initiator and *s_j_* = *k* + 1 for an activated initiator carrying *k* attached barcodes; activation thus sets *s_j_* = 1, and each subsequent attachment increments *s_j_* by unity.

The trajectory is advanced over *S*_steps_ in discrete increments. At step *i*, the number of activation events based on the reaction scheme (1), is drawn from a Poisson distribution with mean *κ_a_I_i_T_i_*,

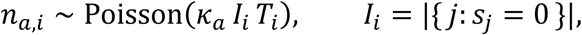

and *n_a_*_,*i*_ inactive initiators, selected uniformly at random, are activated. The target count is updated by

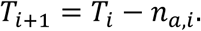

The number of barcode-attachment events in the same step is likewise Poisson-distributed,

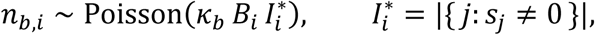

and each event increments the state of a uniformly selected activated initiator. The barcode count is updated by

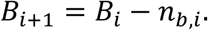

This procedure is the explicit tau-leaping approximation [Reference 23 in main text] to the chemical master equation governing the two reactions, with reaction propensities *a*_1_ = *κ_a_IT* and *a*_2_ = *κ_b_B I*^∗^ evaluated at the current state and a fixed leap *Δt* = 1. It is noted that the accuracy of the approximation depends on the leap condition, namely that the propensities change negligibly over a single step. This assumption could break down as *α* approaches unity.

Upon completion, the distribution of barcodes per initiator is accumulated as a histogram. The inactive initiators and the activated-but-unlabeled initiators are merged into the zero-barcode bin, giving the recorded counts with *k* barcodes, *N*(*k*)

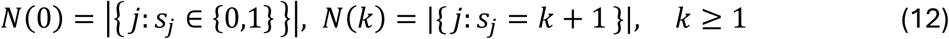

The barcode read-count in Eq. (12) is directly correlated with the quantity *R_i_*(*m*) defined in the “Decoding framework” section of the main manuscript, as the number of reads with *m* matches of the barcode sequence of target *i*. Exact quantitative matches will require additional calibration of the exact number of the different species present in the experiment and its reduced representation in this model. Accordingly, we do not currently emphasize exact quantitative match between theory and experiment. However, we show that the qualitative agreement between the model and experiment is sufficiently high to use this model to guide experimental approach, for example, to explore concentration estimation from the distribution *N*(*k*) as discussed in the “Concentration determination” section of the main text.

#### 5.4 Analytical expression for barcode-count distribution

In addition to the simulation model described in the previous section, we also present an approximate analytical expression for the barcode count distribution *N*(*k*) of Eq.(12) here. The distribution of barcodes among activated initiators is estimated under the assumption that barcode attachment is a memoryless random allocation. Each attachment event is assigned to an activated initiator independently with uniform probability. Under this assumption, the number of barcodes borne by an activated initiator is Poisson-distributed, with a mean equal to the total number of attached barcodes divided by the number of activated initiators, i.e.

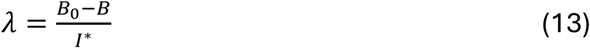

where *B* and *I*^∗^ are evaluated at the end of the trajectory, i.e. after *S_steps_*. The complete initiator population is then a two-component mixture with *I*_0_ − *I*^∗^initiators that were never activated and contribute to the zero-barcode bin, and the *I*^∗^ activated initiators which are Poisson-distributed with mean *λ*. The expected counts are therefore

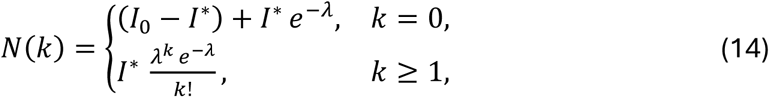

and the probability that a randomly chosen initiator carries *k* barcodes is *P*(*k*) = *N*(*k*)/*I*_0_.

It should be emphasized that the Poisson description is exact only in the limit of many independent allocation events and homogeneous initiator residence times. Deviations are expected where initiators are activated over an extended interval, since initiators that are activated earlier are exposed to barcode attachment for longer than late-activated ones.

#### 5.5 Results

Figure S20 shows the consistency of the continuum ODE model (SI section 5.2) with the stochastic model (SI section 5.3). The number of activated initiators, *I*^∗^, predicted by the stochastic simulation model exactly follows the mean-field ODE Eq. (7) (Figure S20A). Similarly, the number of barcodes, *B*, predicted by the stochastic simulation model follows the mean-field ODE Eq. (9) (Figure S20B). The parameter values used the simulations described in Figure S20 are provided in Supplementary Table 3 below. The excellent quantitative agreement between the stochastic and the ODE models across a range of target concentrations (target fraction, tfrac = *T*_0_/*I*_0_ = 0.1, 0.3, 0.6) provide a basic validation of the theoretical models developed here. The amount of activated initiators at a specific time-point expectedly increases with target concentration as shown in Figure S20A and higher the amount of activated initiators, higher would be the consumption of the barcodes as shown in Figure S20B. Thus the models follow the basic expectations, and the stochastic model can be confidently used to predict observables like the barcode distribution.

**Figure S20:**
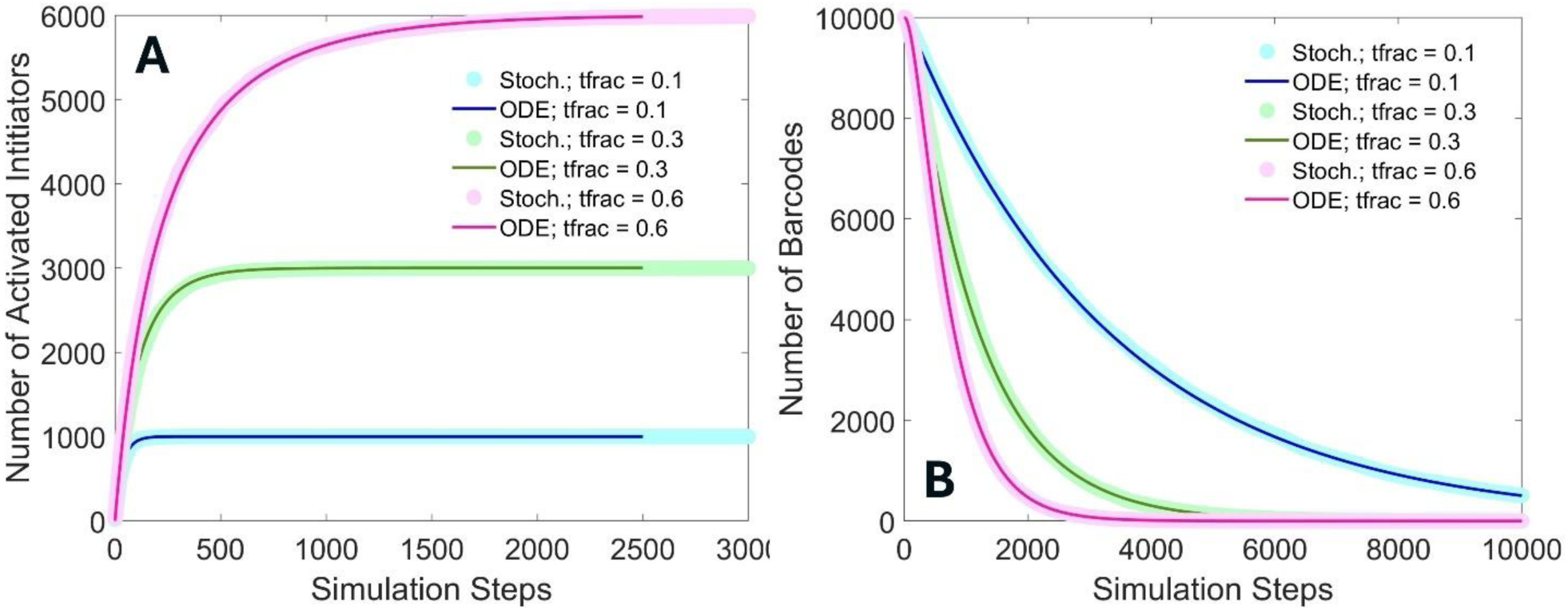
Comparison of Stochastic and Mean-field ODE models. Figure S20A shows the number of activated initiators *I*^∗^ computed using the stochastic simulation method (Stoch. in the legend) to the mean-field model (Eq. 7 in SI section 5.2). Good quantitative agreement is obtained between the two approaches across a range of target concentration fractions (tfrac in the legend, defined as the ratio *T*_0_/*I*_0_). Figure S20B shows the comparison between the two models for the number of Barcodes *B* for the same range of target concentrations. The ODE model is calculated using Eq. 9 in SI section 5.2.

**Supplementary Table 3:** Values of the simulation parameters used to generate Figure S20.

| Symbol | Definition | Value |
| --- | --- | --- |
| $I_0$ | Initial value of initiators | 10,000 |
| $B_0$ | Initial value of barcodes; $B_0 = bfrac * I_0$; $bfrac = 1$ | 10,000 |
| $T_0$ | Initial value of targets; $T_0 = tfrac * I_0$; | variable |
| $\alpha$ | Dimensionless activation reaction rate factor | $3 \times 10^{-3}$ |
| $\beta$ | Dimensionless barcoding reaction rate factor | 1 |
| $k_{max}$ | Maximum number of barcodes considered per strand | 40 |
| $S_{steps}$ | Number of simulation time steps | 10,000 |

Having obtained a reasonable validation of the stochastic model (Figure S20), we proceed to calculate the barcode distribution *N*(*k*), as described in Eq. 12 in section 5.3. We also use Eq. 14 in section 5.4 to understand how effectively the analytical formula captures the results of direct simulations. For proper comparison between the direct simulation and the analytical formula, we calculate the occurrence probability, *P*(*k*) = *N*(*k*)/*I*_0_. The simulation parameter values used are same as those shown in Supplementary Table 3. The results are shown in Figure S21 for target concentration fractions (tfrac defined as the concentration ratio *T*_0_/*I*_0_) ranging from 0.1 to 1.

We see that for low target concentration fractions, tfrac = *T*_0_/*I*_0_, the barcode distribution is approximately unimodal and shifts to a uniformly decreasing function with increasing target concentration fraction. Moreover, the most frequently occurring number of barcode matches decreases as a function of target concentration. For example, the most probable number of barcode matches for tfrac = 0.1 is about 10 matches while for tfrac = 0.6, single barcode matches become the most probable outcome. This is a consequence of the relative magnitudes of the activation reaction rate (*α*) to the barcoding reaction rate (*β*). For the simulations shown in Figure S21, *β* is 300 times larger than *α*. Therefore, for low target concentration fractions, activated initiators are present only in limited amount while there is an excess of barcodes leading to the incorporation of multiple barcodes to the same initiator strand. This effect reduces as the number of activated initiators increases with increasing target fractions causing the most probable barcode numbers to reduce. The other important aspect to take down from Figure S21 is that the analytical model for barcode distribution (Eq. 14 in SI section 5.4) captures the direct simulation results fairly well for low target fractions in the range 0.1 – 0.5, despite its simplicity. However, as the target fraction goes beyond 0.5, there is significant discrepancy between the analytical model and the simulated values. The reason for this deviation is that for low target fractions, the activation reaction reaches saturation quickly and the regime is well described by independent barcode allocation events with stationary activated initiator residence times. This regime is well described by the Poisson distribution. However, when the target fraction is increased the activation events occur over extended times and the condition of a relatively constant activated initiator residence time does not hold across a large fraction of the activated initiators, i.e. initiators that are activated earlier are exposed to barcode attachment for longer than the ones activated later causing the analytical formula to fail. Specifically, it underestimates the effect of the early-activated initiators continuing incorporating barcodes as shown in Figure S21B.

**Figure S21:**
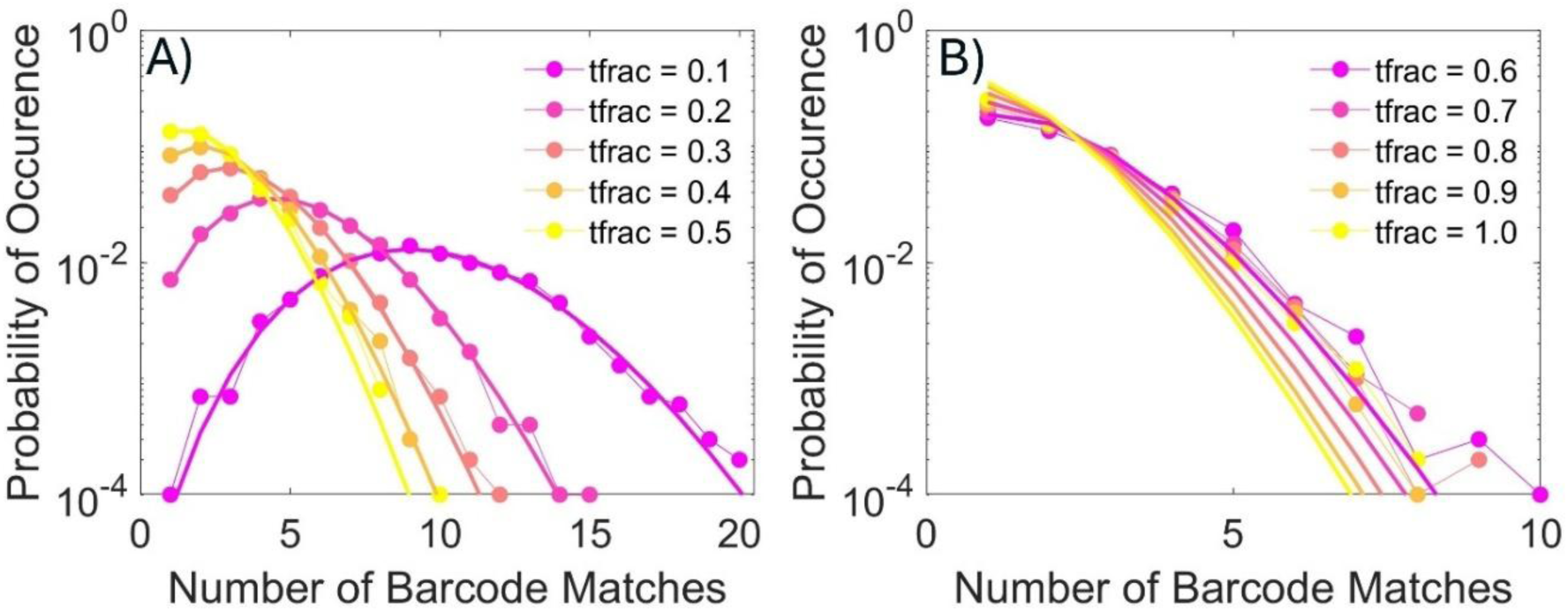
Probability distribution function (PDF) of barcode allotment. A) Shows the probability of occurrence of reads as a function of the number of barcodes incorporated, for example, there is a 1:100 probability that a single read will have 10 barcodes, if the target fraction is 0.1. The analytical formula based on Poisson statistics provides a good fit for the simulated values. B) PDF of barcode incorporation for target concentration fractions higher than 0.5. Here, the Poisson approximation does not hold well and there is significant deviation between the analytical equation and the simulated values.

Figures S20 and S21 are shown for a limited range of parameters for clarity. A more extended parameter sweep, beneficial for a comprehensive sense of system behavior is shown in Figure S22.

**Figure S22:**
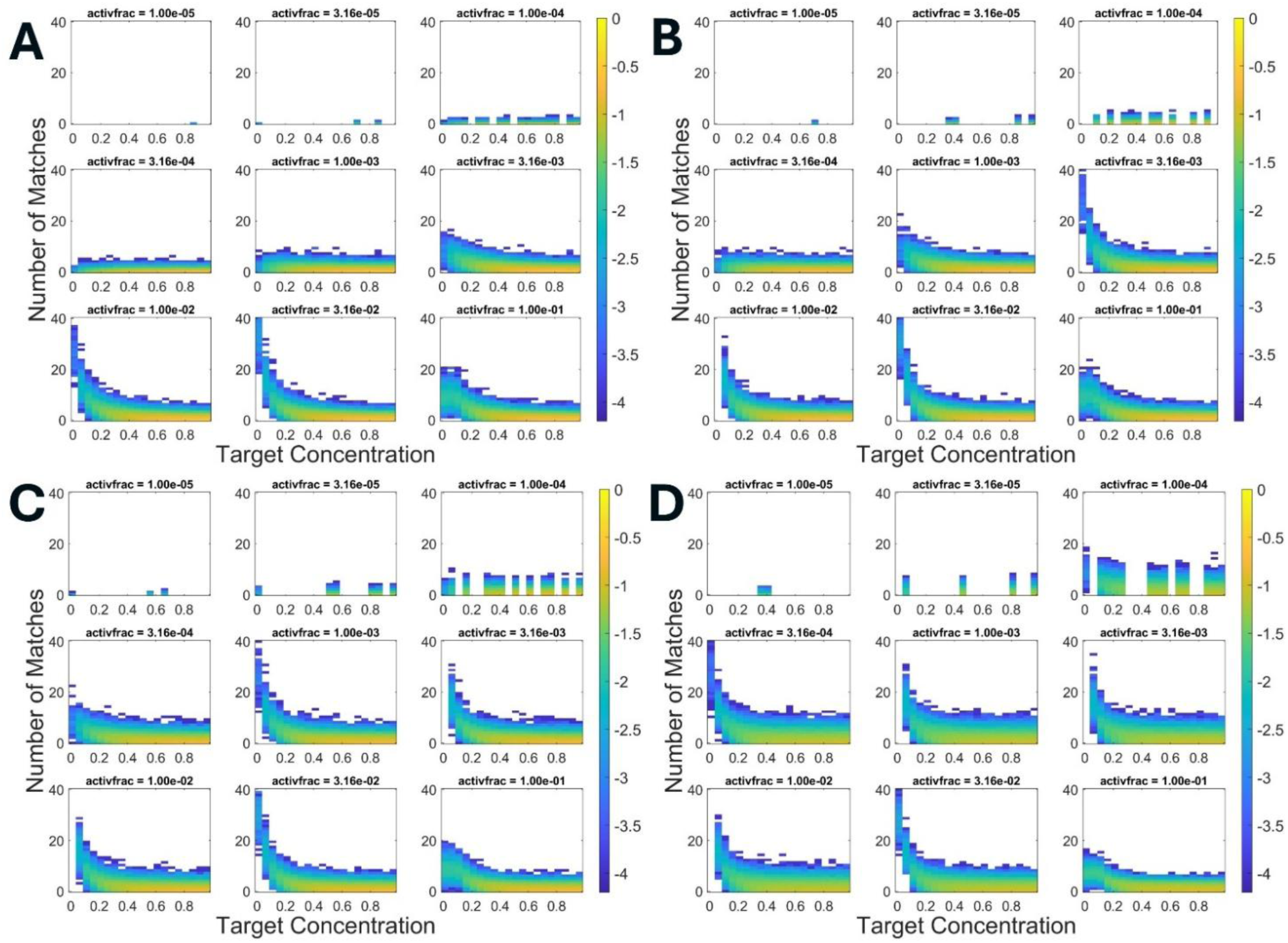
Extended parameter sweep of the stochastic simulation model. The heatmaps presented here show the occurrence probability of the number of barcode matches as a function of target concentration fraction for 0 to 40 matches, for dimensionless barcode reaction rate parameter *β* = 0.3 (Panel A), *β* = 1 (Panel B), *β* = 3 (Panel C), *β* = 10 (Panel D). Within each panel, subpanels correspond to different values of the dimensionless initiator activation reaction rate parameter *α* (activfrac), mentioned in the title of the respective sub-panels.

Inspecting the heatmaps in Figure S22, one can note that for the same activation rate, higher rate of barcoding reaction leads to higher barcode occurrence probability. The barcoding reaction rate is determined by base-pairing free energy of the hairpin sequences *H*_1_ and *H*_2_, whereas the activation reaction rate could originate from aptamer-protein or aptamer-small molecule interaction. The behavior of the occurrence probability is qualitatively similar across a wide range of activation and barcoding reaction rates which presents a challenge for quantification due to insufficient difference in signal. Therefore, at present, the system presented here is best suited to serve as a highly specific multi-class binary (yes/no) assay. Further work is required to develop algorithms to estimate target concentration from measured data, for example by rigorously estimating *α* and *β* parameters for the target of interest. Such directions are open for future studies.

#### 5.6 Non-uniqueness of target concentration determination

To check the effectiveness of the stochastic model we fitted it to the experimentally observed ATP singleplex data. The quantity that was used for this comparison was the probability of occurrence of a read with *k* barcode matches, *P*(*k*). For the simulation *P*(*k*) is obtained by dividing the number of reads with *k* barcode matches, *N*(*k*) from Eq. (12) with *I*_0_. For experimental data *P*(*m*) is obtained by dividing the number of reads with *m* matches, *R*(*m*) defined in the main text, with the total number of reads that were sequenced. Clearly, there can be numerical discrepancy between the two approaches here, but for a first-cut, they appear to be a reasonable choice. For the ATP experiment the initiator concentration *I*_0_ is 1 μM, whereas the target concentration *T*_0_ is 1 mM. However, the reaction can only proceed until either the initiators or the targets are exhausted, therefore the target concentration fraction *T*_0_/*I*_0_ will have a limiting value of 1. The simulation was undertaken with this target concentration fraction. The two free parameters are then the dimensionless rate parameters *α* and *β*. We found a good fit between the experimental data and simulated values for *α* = 2×10^-2^ and *β* = 10 (Main Text Figure 8D, and reproduced here as SI Figure S23A). However, as described in the main text, target concentration is not uniquely determined from sequence data due to its non-monotonic effect on *R*(*m*) (or *N*(*k*)). This is illustrated in Figure S23B below, where a different set of target concentration fraction and rate parameters lead to a close match between the experimental data and simulated values. In this case the simulation parameters are, target concentration fraction of 0.4, *α* = 3×10-3, *β* = 1. The multi-valued behavior implies that determining target concentration from a sample will require a well-calibrated method to link the rate parameters to experimental conditions. Uncertainty in the rate parameters can lead to erroneous or ambiguous concentration determination.

In SI figure S23, we note that the match between the experimental data and simulated values are not good for small values of *m*, say for *m* < 3. As mentioned in the main text, this arises because the matched filters, specifically the read-length filter, removes true matches from the *m* = 1 set. Removing the two filters (SI figure S24A) or removing the read-length filter (SI figure S24B) improves the match between experiment and simulation for small values of *m*. However, for achieving high-fidelity behavior at large *m*, for e.g. *m* > 5 would be preferable where the simulated values are quite close to experimental data irrespective of the number of filters.

**Supplementary Figure S23:**
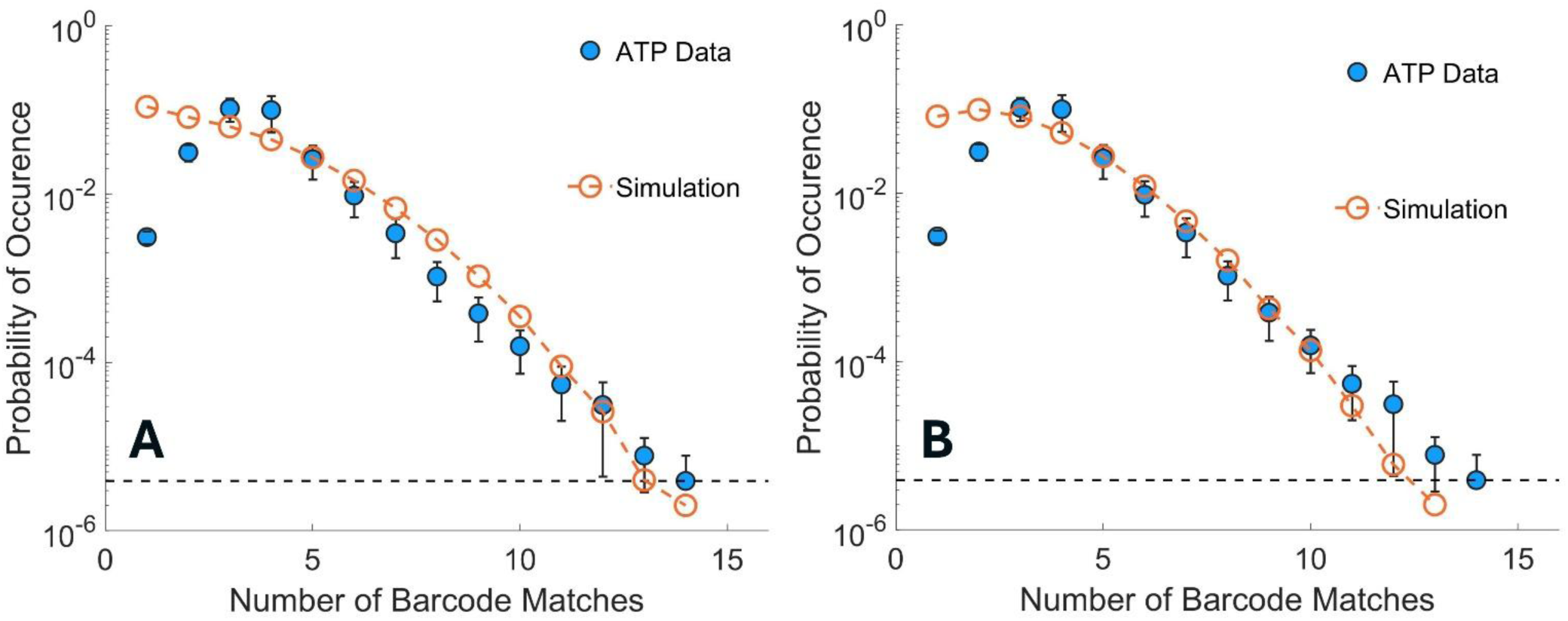
Non-uniqueness of target concentration extracted from sequence data. A) uses target concentration fraction of 1 exactly as in the experiment. B) shows that a different set of parameter values can also lead to a (spurious) match. In this case at target fraction of 0.4, a different set of the dimensionless rate parameters *α* and *β* produce simulated values close to the experimental data.

**Supplementary Figure S24:**
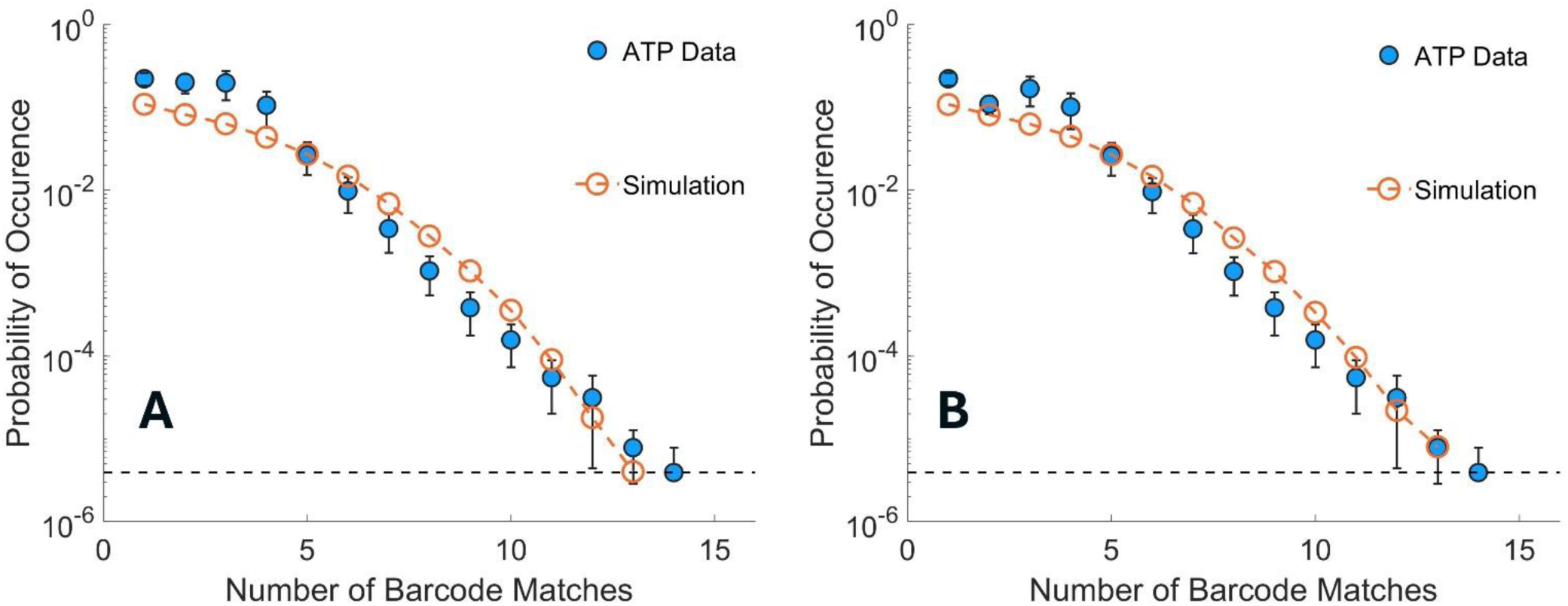
Effect of filters on the match between experimental data and simulated values: A) No-filters applied to the reads, B) only periodicity filter applied. In both cases the match between experimental data and simulated values improve for small number of matches.

